# Oncomodulin derived from regeneration-associated macrophages in dorsal root ganglia after preconditioning injury promotes sensory axon regeneration in the spinal cord

**DOI:** 10.1101/2021.12.28.474322

**Authors:** Min Jung Kwon, Yeojin Seo, Hana Cho, Hyung Soon Kim, Young Joo Oh, Minjae Kim, Hee Hwan Park, Eun-Hye Joe, Myung-Hee Kwon, Han Chang Kang, Byung Gon Kim

**Affiliations:** Department of Brain Science, Ajou University Graduate School of Medicine, Suwon, 16499, Republic of Korea.; Neuroscience Graduate Program, Department of Biomedical Sciences, Ajou University Graduate School of Medicine, Suwon, 16499, Republic of Korea.; Department of Pharmacy, College of Pharmacy, The Catholic University of Korea, Bucheon, 14662, Republic of Korea.; Department of Biomedical Sciences, Graduate School, Ajou University, Suwon 16499, Republic of Korea.; Department of Microbiology, Ajou University School of Medicine, Suwon 16499, Republic of Korea.; Department of Pharmacology, Ajou University School of Medicine, Suwon, 16499, Republic of Korea.; Center for Convergence Research of Neurological Disorders, Ajou University School of Medicine, Suwon, 16499, Republic of Korea.; Department of Neurology, Ajou University School of Medicine, Suwon, 16499, Republic of Korea.

## Abstract

Preconditioning peripheral nerve injury enhances axonal regeneration of dorsal root ganglia (DRG) neurons in part by driving pro-regenerative perineuronal macrophage activation. How these regeneration-associated macrophages influence the neuronal capacity of axon regeneration remains elusive. The present study reports that oncomodulin (ONCM) is an effector molecule derived from regeneration-associated macrophages. ONCM was highly upregulated in DRG macrophages following preconditioning injury and necessary for preconditioning-induced neurite outgrowth. ONCM-deficient macrophages failed to generate neurite outgrowth activity of the conditioned medium in the *in vitro* model of neuron-macrophage interaction. C CL2/CCR2 signaling is an upstream regulator of ONCM since the ONCM upregulation was dependent on CCR2 and CCL2 overexpression-mediated conditioning effects were attenuated in ONCM-deficient mice. Direct application of ONCM potently increased neurite outgrowth in cultured DRG neurons by activating a distinct gene set, particularly neuropeptide-related genes. AAV-mediated overexpression of ONCM construct with the signal sequence increased neuronal secretion of ONCM and enhanced neurite outgrowth in an autocrine manner. For a clinically relevant approach, we developed a nanogel-mediated system for localized delivery of recombinant ONCM to DRG tissue. Electrostatic encapsulation of ONCM by a reducible epsilon-poly(L-lysine)-nanogel (REPL-NG) resulted in a slow release of ONCM allowing sustained bioactivity. Intraganglionic injection of REPL-NG/ONCM complex achieved a remarkable long-range axonal regeneration beyond spinal cord lesion, surpassing the extent expected from the preconditioning effects. NG-mediated ONCM delivery could be exploited as a therapeutic strategy for promoting sensory axon regeneration following spinal cord injury.

## Introduction

Axon regeneration failure following CNS injury is in part due to poor intrinsic capacity of axonal growth in mature CNS neurons (Mar et al., 2014). In contrast, PNS neurons possess the ability of activating axon growth programs upon injury. In a conditioning injury model, preceding injury to the sciatic nerve dramatically enhances the capacity of sensory neurons in the dorsal root ganglia (DRGs) and enables the DRG sensory axons to achieve some degree of re generation following subsequent injury to the spinal cord dorsal column (Richardson and Issa, 1984; Neumann and Woolf, 1999). Many studies attempted to explain how the preconditioning injury can boost intrinsic regeneration potential of DRG neurons, but the precise cellular and molecular mechanisms mediating the conditioning effects remain elusive (Yang et al., 2021). Most of the previous studies have focused on DRG neuronal cells to identify changes in intracellular signaling, transcription factor activation, epigenetic landscape, and metabolic profiles in DRG neurons (Qiu et al., 2002; Jung et al., 2012; Cho et al., 2013; Chandran et al., 201 6; Weng et al., 2017; Rosello-Busquets et al., 2019; Lee et al., 2020). In contrast, relatively little attention was paid to non-neuronal cells surrounding DRG neurons. Recent studies suggest an intriguing possibility that these non-neuronal cells may also contribute to the enhanced regenerative capacity of DRG sensory neurons induced by the preconditioning injury (Poplawski et al., 2018; Avraham et al., 2020; Kalinski et al., 2020; Avraham et al., 2021; Yang et al., 2021).

Several studies reported that peripheral nerve injury leads to activation of macrophage s in DRGs and that the perineuronal macrophages in turn can contribute to the increase in the capacity of axonal growth in DRG sensory neurons (Kwon et al., 2013; Niemi et al., 2013; Zigmond and Echevarria, 2019). Axotomized DRG neurons and surrounding macrophages are highly likely to interact with each other to transmit injury signals from neurons to macrophages and to provide factors to stimulate axon regeneration from the regeneration-associated macrophages to neurons (Kwon et al., 2016; Niemi et al., 2016). The neuron-macrophage interact ion was recapitulated in an *in vitro* system in which conditioned medium obtained from neuron-macrophage co-cultures treated with cAMP provides potent neurite outgrowth activities (Kwon et al., 2013). Employing this neuron-macrophage interaction model, we previously demonstrated that preconditioned DRG neurons generate CCL2/CCR2 chemokine signaling to recruit and activate perineuronal macrophages (Kwon et al., 2015; Kwon et al., 2016). However, it remains to be studied how these regeneration-associated macrophages exert their effects on neurons in return to activate axon growth programs.

Oncomodulin (ONCM) was identified as a myeloid cell-derived molecule that support s regeneration of retinal ganglion neurons following optic nerve injury (Yin et al., 2006; Kurimoto et al., 2013). However, the ability of ONCM to promote axonal regeneration outside of the retina has not been reported. Here, we identify ONCM as an effector molecule derived from regeneration-associated macrophages that are activated by the neuron-macrophage interaction. ONCM expression is increased in DRG macrophages following preconditioning injury in a CCR2-dependent manner. The ONCM upregulation is required for the *in vitro* neuron-macrophage interaction and preconditioning-induced enhancement of axon growth capacity. Furthermore, we demonstrate that delivery of ONCM using a nano-scale hydrogel system, which was recently invented for efficient intracellular delivery of various molecules, peptide drugs in particular (Arnfast et al., 2014; Ye et al., 2016; Hashimoto et al., 2018), into DRGs supports a highly robust extent of sensory axon regeneration well beyond the spinal injury site, implicating ONCM-conjugated nanogel as a potential therapeutic strategy to promote axonal regeneration following CNS injury.

## Results

### Expression of ONCM in DRGs is required for preconditioning effects

ONCM gene expression was examined in the lumbar DRGs following preconditioning sciatic nerve injury (SNI). ONCM mRNA level increased at 3 and 7 days post-injury (dpi) by more than 20 folds (Fig. 1A, B). ELISA measurement of DRG lysates also revealed a marked increase in ONCM protein level at 7 dpi (Fig. 1C). There was no detectable increase in ONCM protein at 7 dpi in ONCM-deficient mice. To ascertain that ONCM is expressed exclusively in DRG macrophages, macrophages were FACS-isolated using Cx 3cr 1-GFP transgenic mice. The number of perineuronal GFP positive cells sharply increased following SNI (Supplementary fig. 1A), and all the GFP positive cells were colocalized with Iba-1 immunoreactivity (Fig. 1D). GFP positive cells were readily separated from GFP negative fractions using flow cytometry (Supplementary fig. 1B). MAP2, a neuron-specific gene, was expressed only in GFP negative fraction, and macrophage specific CD68 was expressed exclusively in GFP positive fraction, indicating a successful separation of macrophage fraction. ONCM mRNA expression was detected only in GFP positive fraction from 7 dpi animals (Fig. 1E), indicating that SNI-induced ONCM upregulation in DRGs occurs exclusively in macrophages surrounding DRG neurons.

**Figure 1.**
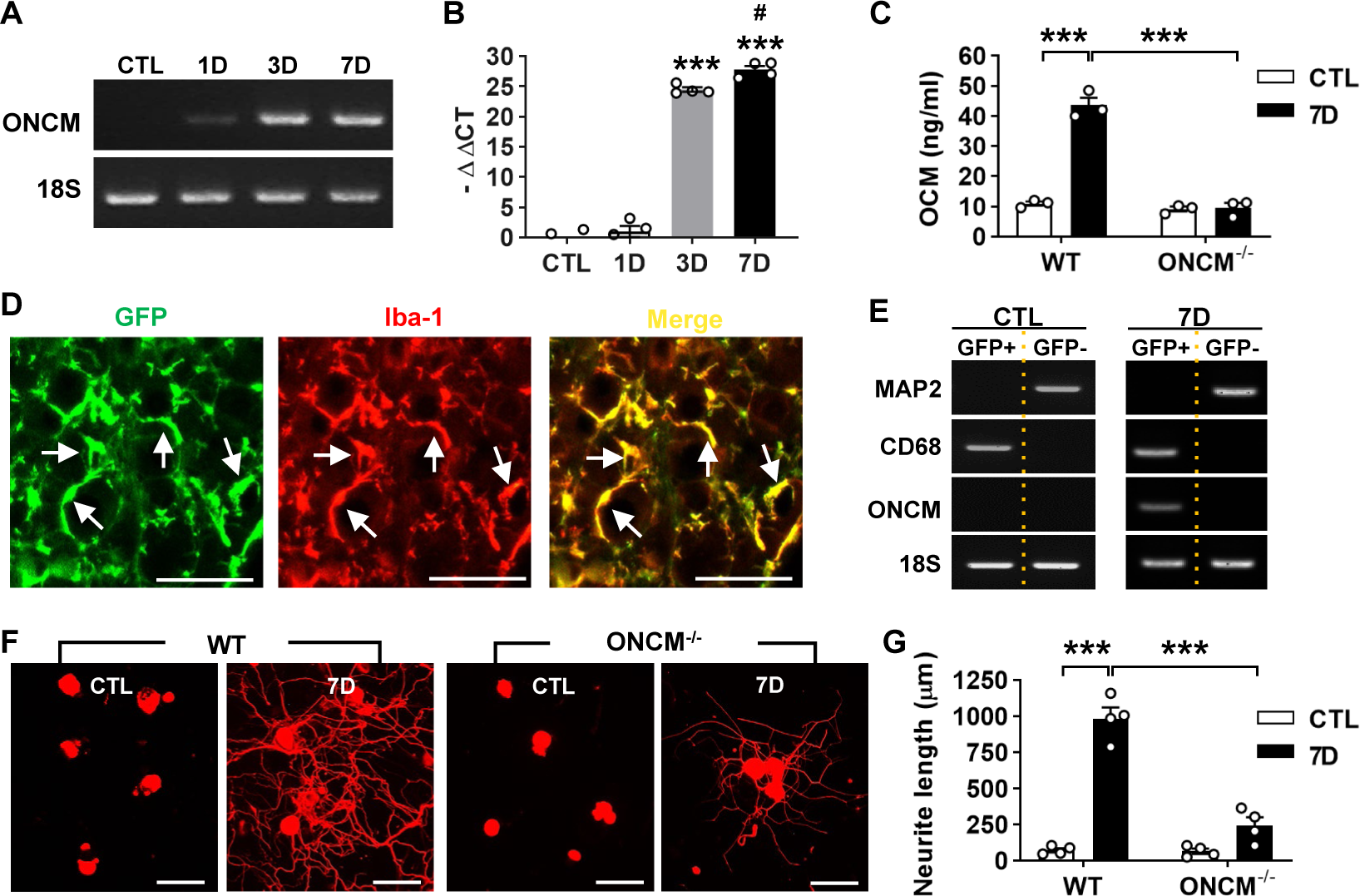
Expression of oncomodulin in DRG macrophages is required for preconditioning effects. A, Representative images of electrophoresed RT-PCR products for oncomodulin (ONCM). 1 8S rRNA was used as an internal reference. L4 and L5 DRGs were obtained 1, 3, and 7 d after ipsilateral sciatic nerve injury (SNI). B, Comparison of ONCM mRNA levels measured by quantitative RT-PCR. N = 4 animals for each time point. \*\*\**p* < 0.001 compared with CTL and 1D values by one-way ANOVA, followed by Tukey’s *post hoc* analysis. #*p* <0.05 compared with 3D values by one-way ANOVA, followed by Tukey’s *post hoc* analysis. C, ELISA measurement of ONCM levels in DRGs from WT and ONCM^-/-^ mice at 0 (CTL) and 7 d after SNI. N = 3 animals for each group. \*\*\**p* < 0.001 compared with CTL values by unpaired *t* test. D, Representative immunofluorescence images of L5 DRG tissue sections obtained from Cx3cr1-GFP mice at 7 d after SNI. Tissue sections were stained with anti-GFP (green) and anti Iba-1 (red) antibodies. Arrows indicate cells positive for both markers. Scale bars represent 100 µm. E, Representative RT-PCR analysis of MAP2, CD68, and ONCM gene expressions from DRG cell fractions separated by FACS. L3, L4, and L5 DRGs from Cx3cr1-GFP mice at 0 (CTL) and 7 d after SNI. MAP2-positive neurons were isolated in the GFP negative fractions. CD68 and ONCM-positive macrophages were collected in the GFP positive fractions. F, Representative images of neurite outgrowth of DRG neurons taken from WT and ONCM^-/-^ m ice 0 (CTL) and 7 d after SNI. DRG neurons from the L3, L4, and L5 DRGs were dissociated and cultured for 15 h before being fixed for the immunofluorescent visualization of neurites with anti-III tubulin. Scale bars represent 100µm. G, Comparison of the mean neurite lengt β h between cultures obtained from WT and ONCM^-/-^ mice at 0 (CTL) and 7 d after SNI. N = 4 animals for each group. \*\*\**p* < 0.001 by unpaired *t* test.

Next, we asked a question whether OCM expression is necessary for the conditioning effects where preconditioning nerve injury enhances capacity of axon growth in DRG sensory neurons (Richardson and Issa, 1984; Neumann and Woolf, 1999). Preconditioning SNI was conducted and the lumbar DRGs were dissected out at 7 dpi for neurite outgrowth assay. The SNI resulted in a huge increase in neurite outgrowth in WT animals (Fig. 1F, G). However, the preconditioning effects on neurite outgrowth were markedly attenuated in ONCM-deficient mice (Fig. 1F, G). Our previous study showed that macrophage activation is essential for maintaining enhanced regenerative capacity, but dispensable for an early induction of the preconditioning effects (Kwon et al., 2013). When lumbar DRG neurons were acutely cultured only at 2 dpi, neurite outgrowth was still increased to an appreciable extent. (Supplementary figure 2A). The enhanced neurite outgrowth activity was not abrogated in ONCM-deficient mice as opposed to a marked reduction of neurite length in ONCM-deficient neurons at 7 dpi (Supplementary Fig. 2A, B), indicating that macrophage-derived ONCM is not involved in inducing preconditioning effects. The intensity of Iba-1 immunoreactivity at the injury site at the sciatic nerve was comparable between WT and ONCM-deficient mice (Supplementary Fig. 3 A). Furthermore, the number of Iba-1 positive perineuronal macrophages and the intensity of Iba-1 immunoreactivity in DGRs were not significantly different between WT and ONC M-deficient mice (Supplementary Fig. 3B), indicating that activation of perineuronal macrophages per se was not affected by ONCM deficiency.

### Macrophage-derived ONCM is necessary for *in vitro* neuron-macrophage interaction

In the *in vitro* neuron-macrophage interaction model, peritoneal macrophages were co-cultured with DRG sensory neurons, and macrophage conditioned medium (MCM) collected fro m the co-cultures with cAMP treatment exhibited robust neurite outgrowth activity (Yun e t al., 2018). In the current study, bone marrow derived macrophages (BMDMs) were used instead of peritoneal macrophages because it is easier for the BMDMs to obtain a large number of macrophages and to maintain stable morphology of cultured macrophages. BMDMs were co-cultured with DRG neurons for 24 h being treated with or without cAMP and then separated from the co-cultured neurons to collect MCM for 72 h thereafter (Fig. 2A). MCM collected from the co-cultured BMDMs with cAMP showed robust neurite outgrowth activity (Fig. 2B, C). However, MCM collected from BMDM only culture treated with cAMP did not show significant neurite outgrowth activity (Fig. 2B, C), confirming that neuron-macrophage interaction is required to generate regeneration-associated macrophages. ONCM concentration in MCM collected from BMDMs co-cultured with neurons accompanied by cAMP treatment was almost 10-fold higher than that in MCM from the co-cultures with PBS (Fig. 2D). ONCM concentration was not significantly elevated in MCM obtained from macrophage only cultures. Next, we examined whether macrophage-derived ONCM is critical in this neuron-macrophage interaction model. When DRG neurons from ONCM deficient mice were co-cultured with wild type (WT) macrophages, MCM from cAMP-treated condition contained neurite outgrowth activity as robust as MCM from all WT cells (Fig. 2E, G). ONCM concentration in this condition was comparable to that in MCM collected from the co-cultures consisting of WT neurons and macrophages (Fig. 2H). When WT DRG neurons were co-culture d with ONCM-deficient macrophages, the neurite outgrowth activity in MCM from cAMP-treated condition was markedly attenuated (Fig. 2F, G). This decrease was associated with a marked decrease in ONCM concentration (Fig. 2H), demonstrating that ONCM derived from re generation-associated macrophages plays an important role for the robust neurite outgrowth activity in the *in vitro* neuron-macrophage interaction model.

**Figure 2.**
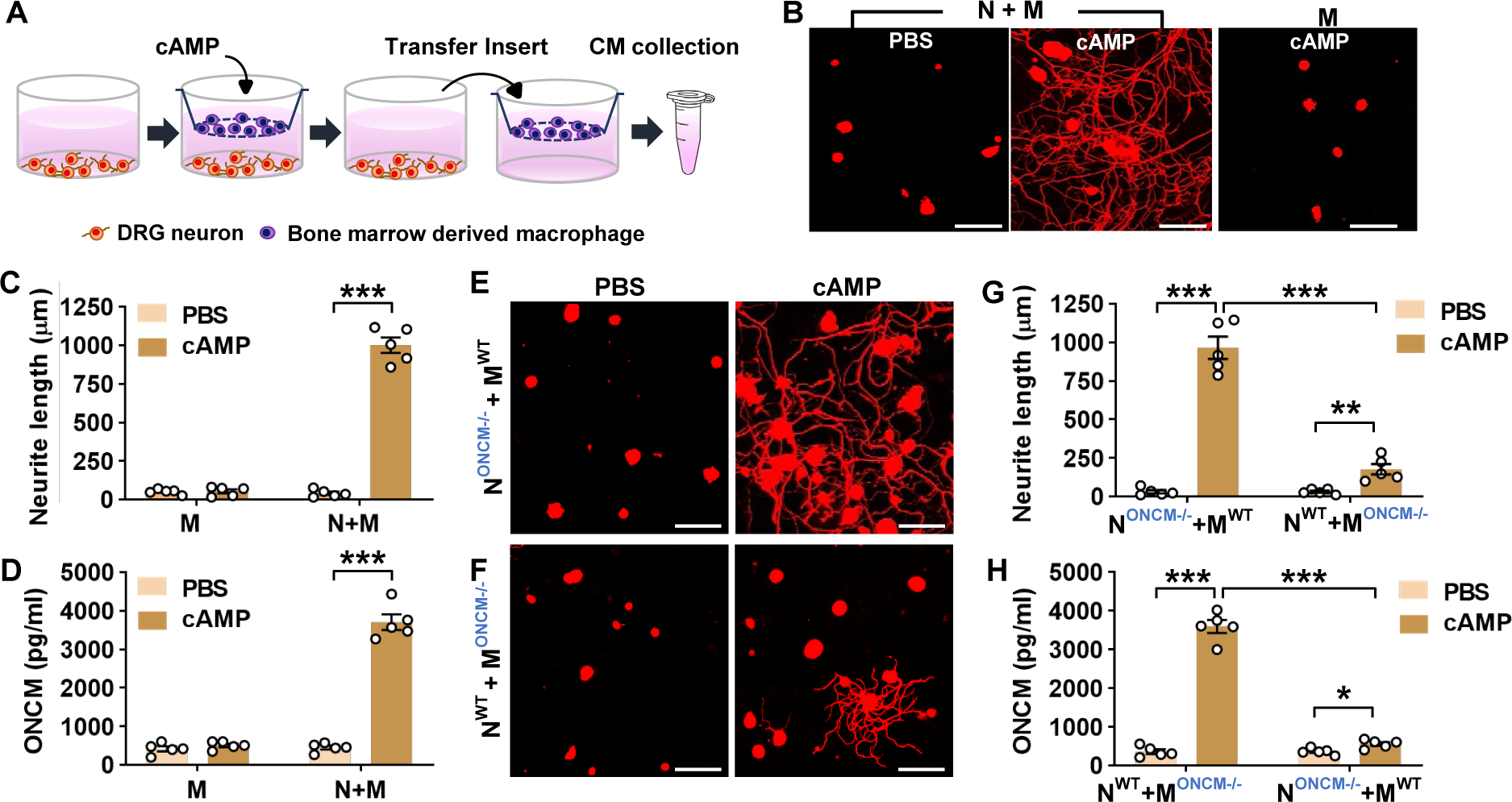
Macrophage-derived ONCM is necessary for *in vitro* neuron-macrophage interaction. A, Schematic diagram illustrating procedures to obtain macrophage conditioned medium (MCM) containing neurite outgrowth activity. Adult DRG neurons (1 x 10^6^ cells per well) are cultured for 4 h, and bone-marrow derived macrophages (5 x 10^6^ cells per well) are cultured on a cell culture insert. At 4 h after plating, either PBS or dibutyryl cAMP is treated. After 24 h incubation, the insert is transferred to another well containing about 1 ml fresh culture medium to collect MCM for 72 h. B, Representative results of neurite outgrowth assay using M CM obtained from various conditions. Cultured DRG neurons and their neurites were visualized by immunofluorescence staining for β III tubulin. Scale bars present 100 µm. C, Quantification graph comparing the mean neurite length. N = 4 independent cultures using independent MCMs for each condition. \*\*\**p* < 0.001 by unpaired *t* test. D, ELISA measurement of the O NCM concentration in the MCM obtained from different culture conditions. N+M, Neuron– macrophage co-cultures; M, macrophage-only cultures. N = 4 independent cultures using independent MCMs for each condition. \*\*\**p* < 0.001 c by unpaired *t* test. E, F, Representative images of neurite outgrowth in DRG neuron cultures treated with MCM obtained from neuron–macrophage co-cultures using WT or ONCM-deficient (ONCM^-/-^) neurons (N) or macrophages (M). All co-cultures were treated with either PBS or db-cAMP (cAMP), and the CMs were collected for 72 h. DRG neurons and their neurites were visualized by immunofluorescence staining for β III tubulin. Scale bars represent 100 µm. G, Quantification graph of neurite out growth in the presence of MCM from cultures of the different neuron–macrophage genotype combinations treated with either PBS or cAMP. N = 4 independent cultures using independent MCMs for each condition. \*\*\**p* < 0.001 and \*\**p* < 0.01 by unpaired *t* test. H, ELISA measurement of the ONCM concentration in the cell culture media obtained from different neuron– macrophage genotype combinations treated with PBS or cAMP. N = 4 independent cultures u sing independent MCMs for each condition. \*\*\**p* < 0.001 and \**p* < 0.05 by unpaired *t* test.

### CCL2/CCR2 chemokine signaling regulates ONCM expression in macrophages

We have previously shown that in the neuron-macrophage interaction model, CCR2-deficient peritoneal macrophages failed to produce pro-regenerative MCM (Kwon et al., 2015). We confirmed that MCM obtained from CCR2-deficent BMDMs co-cultured with WT neurons and treated with cAMP showed neurite outgrowth activity that was markedly attenuated compared to that from WT macrophages (Fig. 3A, B). The reduction of neurite outgrowth activity was accompanied by low levels of ONCM in MCM form CCR2-deficient mice (Fig. 3C). To determine if ONCM production *in vivo* is influenced by CCL2/CCR2 signaling, we performed preconditioning SNI in CCR2-deficient mice and measured the level of ONCM in DRGs. The level of ONCM concentration in DRG tissue increased at 7 dpi regardless of genotype (Fig. 3D). However, the extent of SNI-induced increase in the level of ONCM was significantly smaller in CCR2-deficient mice than that in WT animals.

**Figure 3.**
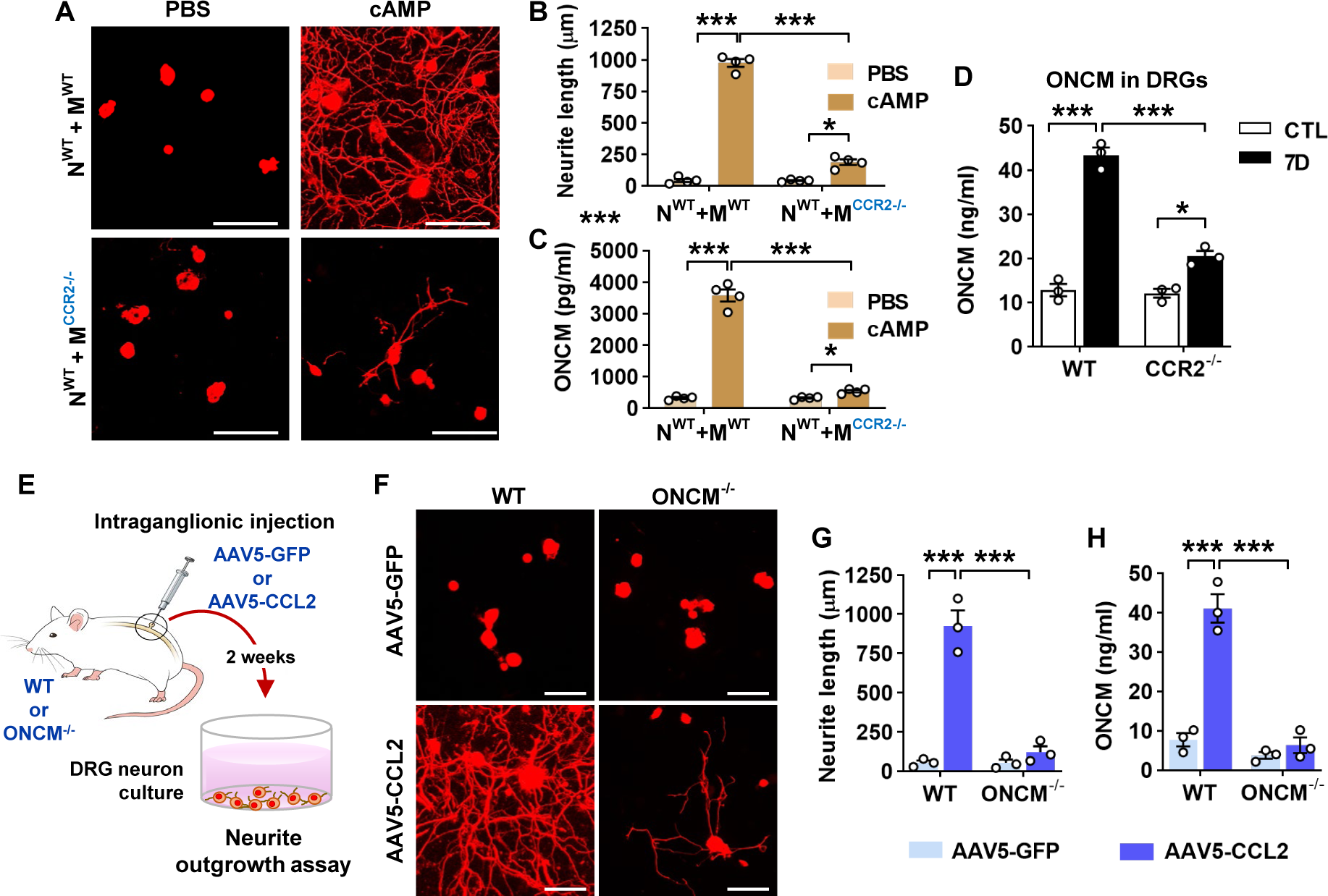
CCL2/CCR2 chemokine signaling regulates ONCM expression in macrophage s. A, Representative images of neurite outgrowth in DRG neuron cultures treated with macroph age conditioned medium (MCM) obtained from neuron–macrophage cocultures using WT or CCR2-deficient (CCR2^-/-^) neurons (N) or macrophages (M). All cocultures were treated with either PBS or db-cAMP (cAMP), and the MCMs were collected for 72 h. DRG neurons and t heir neurites were visualized by immunofluorescence staining for β III tubulin. Scale bars rep resent 100 µm. B, Quantification graphs of neurite outgrowth in the presence of MCM from cultures of the different neuron–macrophage genotype combinations treated with PBS or cAM P. N = 4 independent cultures using independent MCMs for each condition. \*\*\**p* < 0.001 and \**p* < 0.05 by unpaired *t* test. C, ELISA measurement of the ONCM concentration in the cell culture media obtained from different neuron–macrophage genotype combinations treated with PBS or cAMP. N = 4 independent cultures using independent MCMs for each condition. \*\*\**p* < 0.001 and \**p* < 0.05 by unpaired *t* test. D, ELISA measurement of ONCM levels in DR Gs from WT and CCR2^-/-^ mice 0 (CTL) and 7 d after sciatic nerve injury. *n* = 4 animals for each group. \*\*\**p* < 0.001 and \**p* < 0.05 by unpaired *t* test. E, Experimental schematic diagram depicting the experimental processes. F, Representative images of neurons cultured from L5 DRGs freshly dissected from WT or ONCM^-/-^ mice subjected 2 weeks previously to intraganglionic injection of AAV5-GFP or AAV5-CCL2. The culture period was 15 h and DRG neurons and their neurites were visualized by immunofluorescence staining for β III tubulin. Scale bars represent 100µm. G, H, Quantification graphs comparing the mean neurite length (G) and the protein level of ONCM (H). N = 3 animals for each group. \*\*\**p* < 0.001 and \*\**p* < 0.0 1 by unpaired *t* test.

We have previously shown that intraganglionic injection of AAV5-CCL2 induced perineuronal macrophage activation and mimicked conditioning effects on neurite outgrowth in rats (Kwon et al., 2015). We performed intraganglionic injection of AAV5-CCL2 into L5 lum bar DRGs in WT or ONCM deficient mice and freshly cultured these DRG neurons 2 weeks after the injection (Fig. 3E). Consistent with the previous study, intraganglionic AAV5-CCL2 injection resulted in marked enhancement of neurite outgrowth in WT mouse DRG neurons (Fig. 3F, G). This increase in neurite outgrowth was accompanied by a huge increase in ONC M levels in DRGs (Fig. 3H). However, the effects of CCL2 overexpression on neurite outgrowth were largely attenuated in ONCM deficient mice (Fig. 3F, G), and the level of ONCM concentration was not affected by CCL2 overexpression in ONCM deficient mice (Fig. 3H). The number of perineuronal Iba-1 positive macrophages and the intensity of Iba-1 immunoreactivity in DRGs following AAV5-CCL2 injection were comparable between WT and ON CM-deficient mice (Supplementary fig. 4A, B), corroborating that macrophage activation by CCL2/CCR2 signaling is not dependent on ONCM.

### ONCM potently increases neurite outgrowth in cultured DRG neurons by upregulating a distinctive set of RAGs

The finding that ONCM is required for the conditioning effects prompted us to test whether a direct application of recombinant ONCM protein to cultured DRG neurons can increase neurite outgrowth. ONCM treatment promoted neurite outgrowth in a dose-dependent manner, with the extent of neurite outgrowth by 100 ng ONCM comparable to that achieved by preconditioning SNI (Fig. 4A, C). Treatment of ONCM also overcame the growth inhibitory influence of CSPGs as coated substrate in a dose-dependent manner (Fig. 4B, D). Thus, ONCM possesses robust neurite outgrowth activity for cultured DRG sensory neurons. We compared the neurite outgrowth activity of ONCM with that of molecules with reported neurite outgrowth activity such as nerve growth factor (NGF) and IL-6 (Cafferty et al., 2001; Cafferty et al., 2004). Treatment of NGF at a dosage of either 10 or 100 ng did not enhance neurite outgrowth (Fig. 4E, G). Hyper IL-6, a fusion protein of IL-6 and soluble IL-6 receptor facilitating intracellular transduction of IL-6 (Fischer et al., 1997), was also not effective. When hyper IL-6 was treated to DRG neuron primed with NGF, the extent of neurite outgrowth was significantly increased (Fig. 4E, G). However, the neurite outgrowth activity of ONCM was still more robust than that of hyper IL-6 for NGF-primed neurons. It was reported that ONCM is effective in promoting optic nerve regeneration when combined with cAMP {Yin, 2006 #1486}. In vitro, ONCM increased neurite extension of retinal ganglion neurons when treated together with mannose and forskolin. In our experiment with DRG sensory neurons, ONCM alone showed very strong neurite outgrowth activity, and combination of cAMP or mannose plus forskolin did not produce any significant additive effects (Fig. 4F, H).

**Figure 4.**
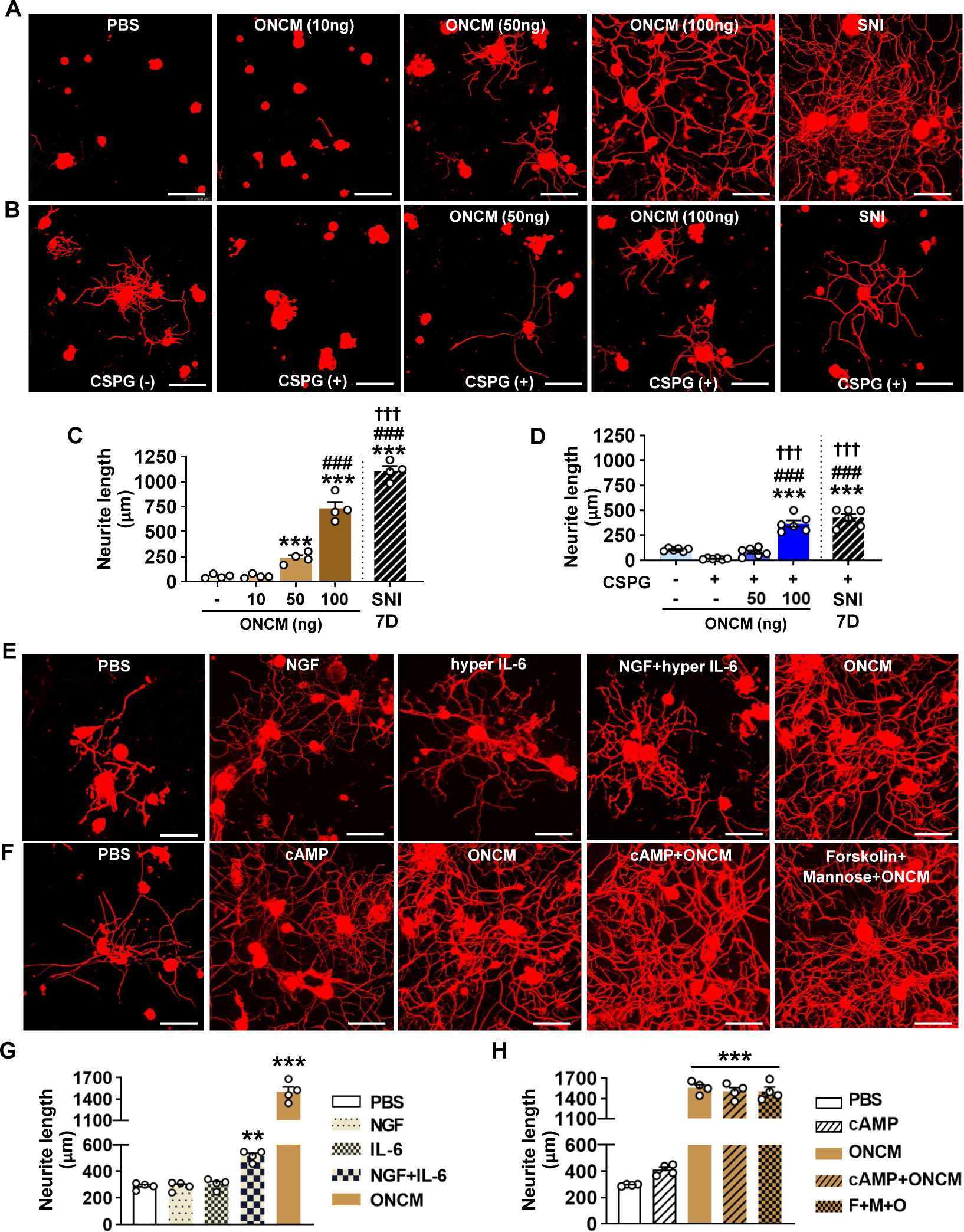
ONCM potently increases neurite outgrowth in cultured DRG neurons. A, Representative images of β III-tubulin-positive cultured DRG neurons. Adult DRG neuron s were cultured for 15 h with PBS or recombinant oncomodulin (ONCM) at 10, 50, and 10 0 ng concentrations. DRGs were freshly dissected from animals at 7 d after sciatic nerve injury (SNI) as positive control. Scale bars, 100µm. B, Representative images of neurite outgrowth of DRG neurons grown on a substrate precoated with chondroitin sulfate proteoglycans (CSPGs) for 24 h in response to different dosages of ONCM. DRG neurons dissected from animals at 7 d after SNI as positive control. At the 15 h culture period, neurite length was not appropriate to compare between the groups. Scale bars represent 100 µm. C, Quantification of t he neurite outgrowth assay with ONCM treatment. N = 4 independent cultures for each condition. *** *p* < 0.001 and * *p* < 0.05 compared with PBS and 10 ng ONCM groups, ### *p* < 0.001 compared with 50 ng ONCM group, and ††† *p* < 0.001 compared with 100 ng ONCM treatment group by one-way ANOVA followed by Tukey’s *post hoc* analysis. D, Quantification graph of the neurite outgrowth on CSPG coated substrate. pre-coated CSPG o n neurite outgrowth assay with recombinant oncomodulin treatment. N = 5 independent cultures for each condition. \*\*\**p* < 0.001 compared with CTL value, ###*p* < 0.001 compared with CSPG alone group, and ¶¶¶ *p* < 0.001 compared with CSPG with 50 ng ONCM treatment group by one-way ANOVA followed by Tukey’s *post hoc* analysis. E, Representative images of β III tubulin-positive cultured DRG neurons grown for 24 h with PBS, NGF, hyper IL-6, hyper IL-6 with NGF priming, and ONCM treatments. Scale bars represent 100 µm. F, Representative images of neurite outgrowth in DRG neuron cultures treated with PBS, cAMP, ONCM, combination of cAMP and ONCM, and combination of mannose, forskolin, and ONCM. Scale bars represent 100 µm. G, H, Quantification graphs of neurite outgrowth of cultured DRG neurons. N = 4 independent cultures for each condition. \*\*\**p* < 0.001 and \*\**p* < 0.01 by one-way ANOVA followed by Tukey’s *post hoc* analysis.

In order to gain insight on potential mechanisms downstream of ONCM for this potent neurite outgrowth activity, we performed RNA-seq using cultured DRG neurons treated with ONCM for 12 h. Since the method of culturing DRG neurons cannot completely remove no n-neuronal cells, we enriched neurons using bovine serum albumin (BSA) cushion (Weng et al., 2016). BSA cushion effectively diminished expression of several non-neural cell-specific genes (Supplementary fig. 5). Transcriptomic analysis revealed that 1321 and 433 genes were upregulated and downregulated, respectively, by more than 2 folds (Fig. 5A). It was notable that several neuropeptide family genes such *Npy* (Neuropeptide Y), *Gal* (Galanin), and *Vip* (Vasoactive intestinal peptide) were highly upregulated. Gene set enrichment analysis (GSEA) showed that the gene set related to neuropeptide hormone activity was highly enriched in the upregulated genes in response to ONCM treatment (Fig. 5B). GO analysis also revealed that neuropeptide signaling terms are enriched (Fig. 5C). Upregulation of the neuro peptide genes were validated in an independent DRG neuron sample (Supplementary fig. 6A). Furthermore, we found that expression of *Npy* and *Gal*, along with *Sprr1A,* which is a well characterized RAG and highly upregulated in the transcriptomic analysis, were highly induced in DRGs following SNI (Supplementary fig. 6B). The increases of those genes at 7 dpi were markedly attenuated in ONCM deficient mice.

**Figure 5.**
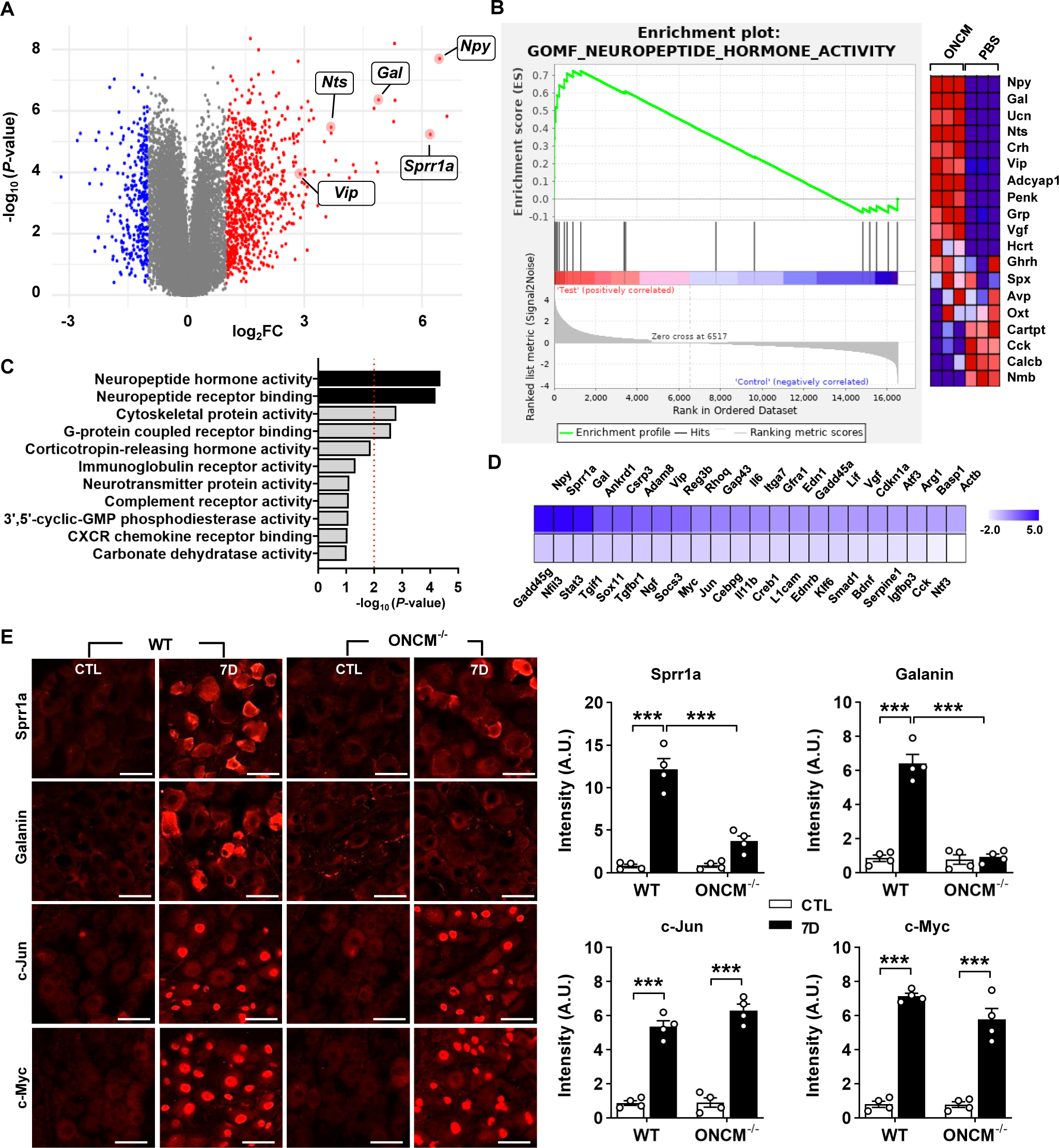
ONCM upregulates a distinctive set of regeneration-associated genes. A, Volcano plot of the gene expression profiles in DRG neurons treated with oncomodulin (O NCM). Blue and red dots indicate genes revealing a < 1.5-fold decrease and a > 1.5-fold increase in expression by ONCM, respectively. N = 3 biological replicates (independent cultures) f or each group. B, Gene set enrichment analysis (GSEA) of ONCM-upregulated genes. The G SEA algorithm calculated an enrichment score reflecting the degree of over representation at the top or bottom of the ranked list of the genes included in a gene set in a ranked list of all genes present in the RNA-seq dataset. The analysis demonstrates that genes related to neuropeptide hormone activity are enriched in the ONCM treatment group. C, Gene ontology analysis of upregulation genes revealing a > 1.5-fold increase in expression by ONCM. Top 11 GO terms for molecular function ranked by fold enrichment were shown in the bar graph. Black bars indicate neuropeptide-related GO term. E, Color-coded heatmap of the gene expression levels for selected 44 RAGs. F, Representative images of Sprr1a, Galanin, c-Jun, and c-Myc immunoreactivity in DRGSs of WT and ONCM^-/-^ mice at 0 (CTL) and 7 d after SNI. Quantification of intensity of immunoreacitivities in WT and ONCM^-/-^ mice 0 (CTL) and 7 d after SNI. N = 4 animals for each condition. Scale bars represent 50 μm. \*\*\**p* < 0.001 by one-way ANOVA followed by Tukey’s *post hoc* analysis.

Preconditioning SNI elicits upregulation of a group of genes that are collectively called “regeneration-associated genes (RAGs)”(Ma and Willis, 2015; Chandran et al., 2016). We previously selected a set of 44 RAGs that had been consistently reported as upregulated genes in previously published gene expression studies in DRGs (Shin et al., 2020). To determine potential contribution of ONCM to the SNI-induced RAG profile, we analyzed the expression of the 44 RAGs in ONCM-treated cultured DRGs. Most of the neuropeptide-related genes upregulated by ONCM (such as *Npy*, *Gal*, and *Vip*) belonged to the selected RAG set (Fig. 5D). *Sprr1a* and *Gap43*, which are representative genes reflecting the intrinsic capacity of axon growth, were significantly affected by ONCM. ONCM also significantly increased expression of *Ankrd1*, *Csrp3* (also known as *Mlp*), and *Reg3b*, of which neurite outgrowth activity was experimentally demonstrated (Tam et al., 2002; Stam et al., 2007; Levin et al., 2019). RAGs related to ECM and cell adhesion (*Adam8* and *Itga7*) and cytokine/growth factors (*Il6*, *En1*, *Gfra1*) were also significantly, but to a lesser extent, increased in response to ONCM. In contrast, expression of transcription factor RAGs such as *Stat3*, *Myc*, *Jun*, and *Smad2*, which are known to play a role of hub transcription factors to regulate modular expression of RAG subs ets (Chandran et al., 2016; Shin et al., 2020), was not significantly changed (Fig. 5D). To ascertain whether certain RAGs are independent of ONCM, we compared the protein expression of RAGs following SNI between wild type and ONCM-deficient mice. As expected, immunoreactivities against Galanin and Sprr1A were dramatically induced at 7 days following SN I and noticeably attenuated in ONCM-deficient mice (Fig. 5E, F). However, the injury-induced nuclear immunoreactivities to c-Jun and c-Myc transcription factors were not reduced in ONCM-deficient mice (Fig. 5E, F).

### Overexpression of ONCM construct with the signal sequence in DRGs enhances axon growth capacity

We sought to overexpress ONCM in DRGs to examine whether ONCM is sufficient to enhance capacity of axon regeneration in DRG sensory neurons. When ONCM plasmids were transfected into HEK 293 cells, ONCM expression was detected in cell lysates but not in the culture medium (Fig. 6A). ONCM cDNA was also delivered to Raw 264.7 mouse macrophage cells, but still the level of ONCM secreted to the media was negligible (Fig. 6A), suggesting that exogenously delivered ONCM gene could not utilize the secretory machinery in macro phages. We found that ONCM gene is lacking the signal sequence at its N-terminus, which encodes the signal peptide that directs newly synthetized proteins toward the secretory pathway. To facilitate extracellular secretion of ONCM, DNA sequences encoding bacterial alkaline phosphatase signal peptide were inserted into ONCM cDNA construct (Fig. 6B). Transfection of the ONCM construct with the signal sequence (SS-ONCM) immensely increased the amount of ONCM in the culture medium from HEK 293 and mouse macrophage cells (Fig. 6 A). Since virus-mediated gene delivery to macrophages *in vivo* is challenging (Zhang et al., 2 009), we conceived an alternative approach of AAV-mediated transfection of DRG neurons with ONCM construct containing the signal sequence, delivering ONCM to the neurons in an autocrine manner (Fig. 6C). We confirmed that adjoining the signal sequence greatly improved ONCM secretion in PC12 neuronal cells as well (Fig. 6D). Intraganglionic injection of AAV5-ONCM resulted in a slight but significant increase in neurite length when the DRGs were acutely cultured (Fig. 6E, F). In comparison, AAV5-SS-ONCM containing the signal sequence dramatically increased the extent of neurite outgrowth by almost 10 folds, supporting an idea that extracellularly secreted ONCM is sufficient to increase the capacity of axon growth in DRG sensory neurons.

**Figure 6.**
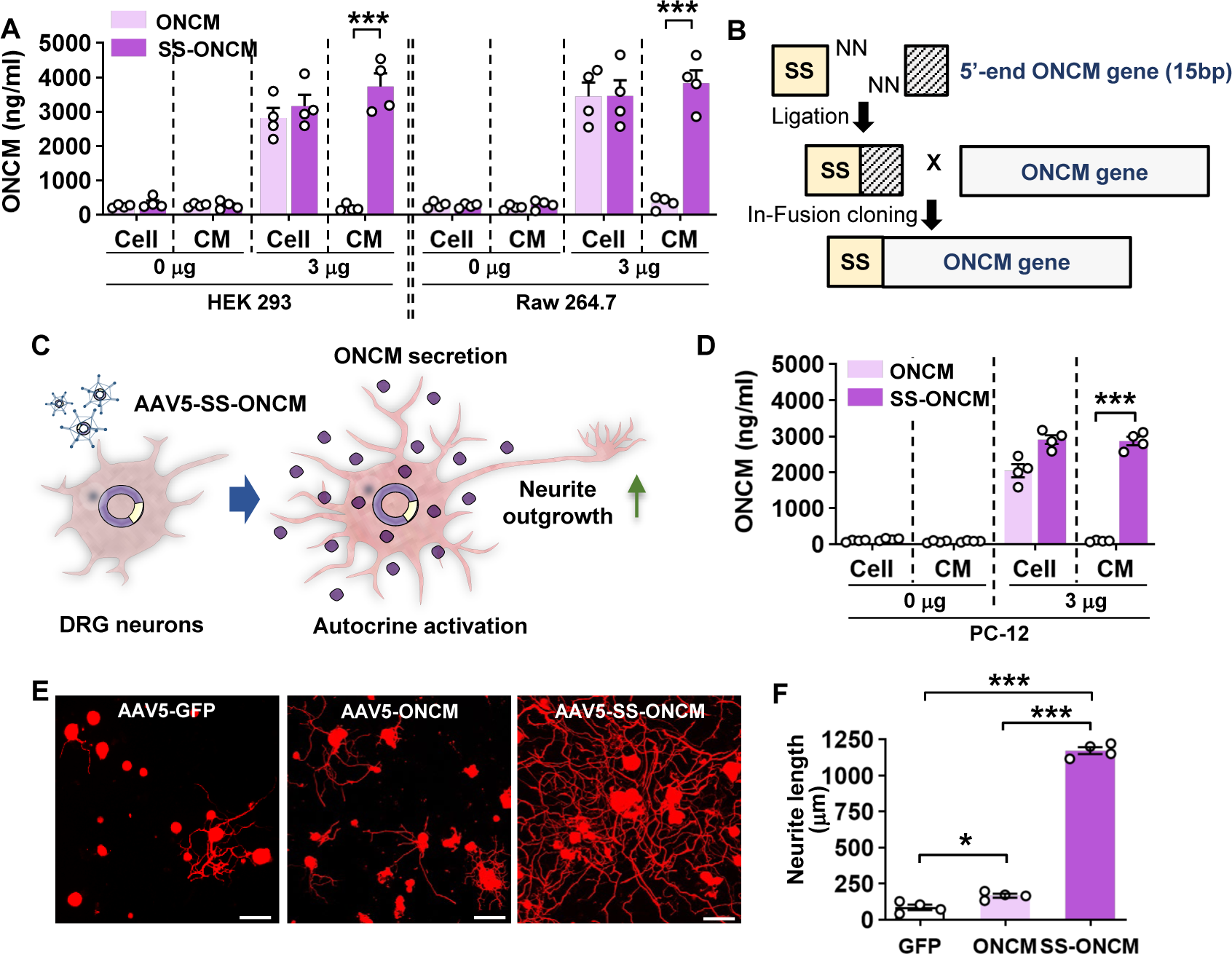
Overexpression of oncomodulin construct with the signal sequence in DRGs enhances axon growth capacity. A, ELISA results for the oncomodulin (ONCM) level in the cell lysates and the culture medium from cultured HEK 293 and Raw 264.7 cells transfected with ONCM with or without the signal sequence (SS). N = 4 independent cultures for each condition. \*\*\**p* < 0.001 by unpaired *t* test. B, Schematic representation of a construct with the signal sequence (SS) for secretion of ONCM. C, Schematic diagram of AAV-mediated transfection with ONCM construct containing the signal sequence (AAV5-SS-ONCM) in the DRG neurons and autocrine activation of neurite outgrowth. D, ELISA measurement of the ONCM concentration in the cell lysate a nd the conditioned medium from cultured PC 12cells transfected with ONCM with or without the signal sequence (SS). N = 4 independent cultures for each condition. \*\*\**p* < 0.001 by unpaired *t* test. E, Representative images of neurons cultured from L5 DRGs freshly dissected from animals subjected to intraganglionic injection of AAV5-GFP, AAV5-ONCM or AAV5-S S-ONCM 2 weeks prior to culture. The culture period was 15 h and DRG neurons and their neurites were visualized by immunofluorescence staining for β III tubulin. Scale bars represent 100 µm. F, Quantification graph comparing the mean neurite length. \*\*\**p* < 0.001 and \**p* < 0.01 by one-way ANOVA followed by Tukey’s *post hoc* analysis.

### Encapsulation via a nanogel drug delivery system sustains stability of ONCM activity

To develop an ONCM-based therapeutic strategy that is more amenable to clinical translation, we attempted to deliver recombinant ONCM protein to DRGs. We tested whether recombinant ONCM injection to DRG neurons could enhance the extent of neurite outgrowth from DR G neurons that were acutely cultured at 3, 7, and 14 dpi. Injection of the recombinant ONC M protein increased the mean neurite length at 7 and 14 dpi, but the extent of neurite outgrowth was much less that what is expected from preconditioning SNI (Supplementary fig. 7A, B).

The apparent difference between the ONCM effects *in vitro* and *in vivo* led us to consider an issue of ONCM protein stability when delivered to the DRG tissue *in vivo*. To test if biological activity of ONCM would be affected by *in vivo* tissue environment, we added fetal bovine serum at different concentrations to the culture medium where DRG neurons were cultivated with ONCM. Surprisingly, addition of the serum attenuated the neurite outgrowth activity of ONCM in a dose-dependent manner (Fig. 7A, B), suggesting that ONCM activity would be strongly inhibited by tissue fluids in an *in vivo* environment. To improve the stability of ONCM protein *in vivo*, we developed a novel nanogel-based drug delivery system, reducible c-poly(_L_-lysine) (EPL)-based nanogel (REPL-NG). Positive charges of the primary amines in REPL-NG could encapsulate ONCM, which has a strong negative charge (pI = 3.9), via an electrostatic force and thereby increase its *in vivo* stability (Fig. 7C). When nanogel-encapsulated ONCM was treated instead of recombinant ONCM, neurite outgrowth was robustly enhanced regardless of adding 10% serum (Fig. 7D, E), demonstrating that R EPL-NG/ONCM complex successfully protected ONCM activity from the serum components.

**Figure 7.**
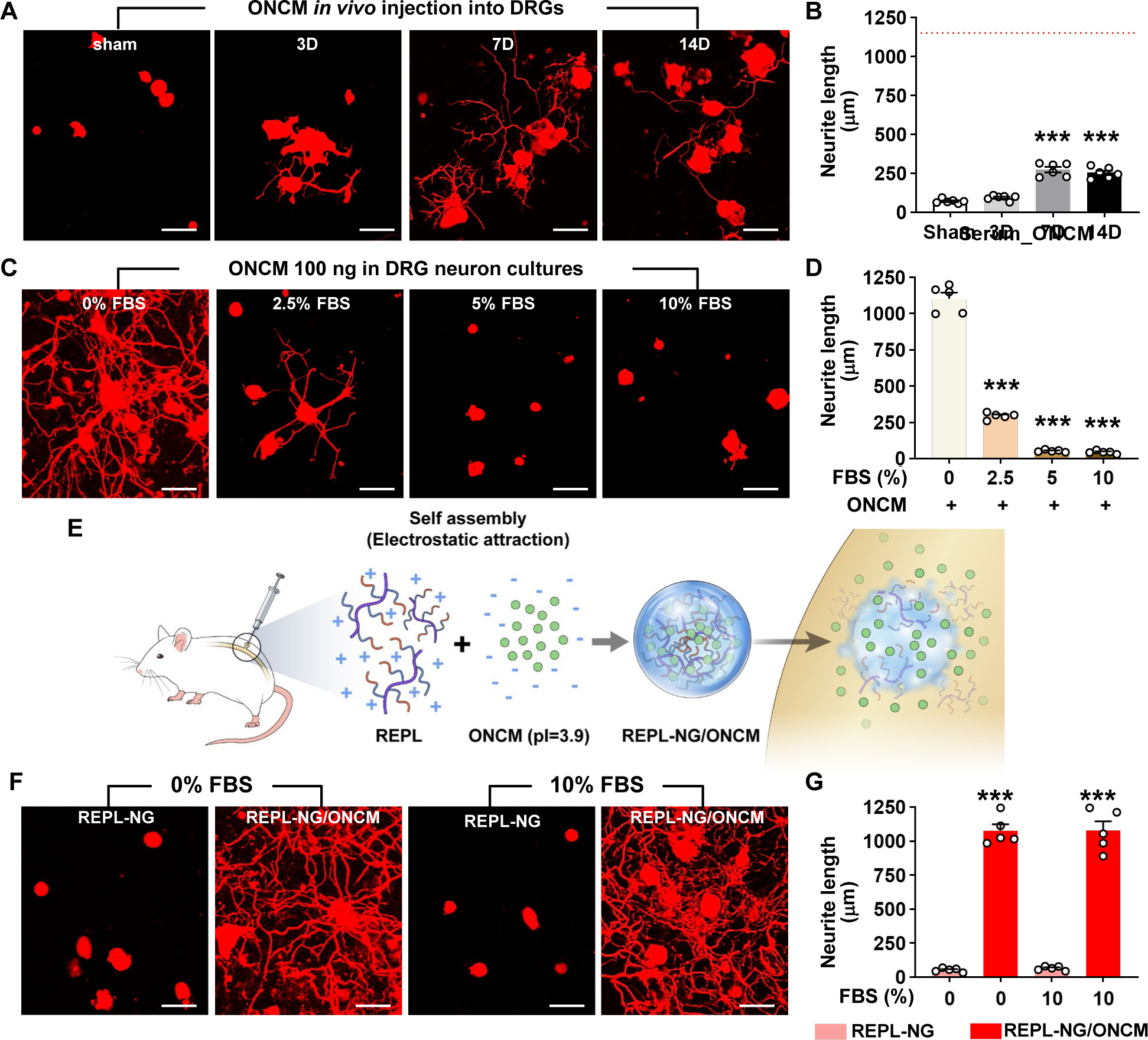
Encapsulation via a nanogel drug delivery system sustains stability of ONCM activity. A, B, Comparison of the mean neurite length between cultures from intraganglionic injection of ONCM 0 (sham), 3, 7, and 14 d after injection. Scale bars, 100mm. N = 6 animals per group. \*\*\**p* < 0.001 compared with control values by one-way ANOVA followed by Tukey’s *pos t hoc* analysis. C, Adult DRG neurons were cultured for 15 h with FBS at 0, 2.5, 5, and 10% c oncentrations treated with ONCM. Scale bars, 100mm. D, Quantification of the DRG neurite outgrowth assay. N = 5 independent cultures for each condition. \*\*\**p* < 0.001 compared with 0% FBS with ONCM value by one-way ANOVA followed by Tukey’s *post hoc* analysis. E, A simplified illustration of the *in vivo* ONCM delivery system by nanogel-based drug deliver y system, reducible c-poly(_L_-lysine) (EPL)-based nanogel (REPL-NG). F, Neurite outgrowth of cultured DRGs without FBS or 10% FBS treated with REPL-NG or REPL-NG/ONCM. T he culture duration was 15 h. Scale bars, 100mm. G, Quantification of the DRG neurite outgr owth assay. N = 5 independent cultures for each condition. \*\*\**p* < 0.001 compared with 10% FBS with REPL-NG value by one-way ANOVA followed by Tukey’s *post hoc* analysis.

The REPL-NG/ONCM complex was injected into the DRGs to examine the extent of neurite outgrowth from acutely cultured DRGs. REPL-NG with ONCM conjugated at 5:1 ratio, REPL-NG/ONCM (5:1), significantly enhanced axon growth of DRG neurons compared to no injection control at 14 dpi (Supplementary fig. 7C). However, the extent of growth was still smaller than that achieved by preconditioning SNI (Supplementary fig. 7C, D). Then, we increased the composition of ONCM to generate REPL-NG/ONCM (3:1) that can deliver a higher amount of ONCM. Injection of the REPL-NG/ONCM (3:1) increased neurite outgrowth at 14 dpi more markedly than the REPL-NG/ONCM (5:1), to the extent comparable to that achieved by preconditioning SNI (Fig. 7F, G). Injection of REPL-NG without ONCM did not affect neurite outgrowth. Encapsulation of ONCM by REPL-NG is supposed to result in a slow release of free ONCM. To test this notion *in vivo*, we tried to measure the level of released ONCM in DRG tissue following injection of the nanocomplex. We first confirmed that ELISA cannot detect encapsulated ONCM (REPL-NG/ONC M) even though it was exposed to tissue lysis buffer containing detergent (Supplementary fig. 8A, B). REPL-NG possesses multiple disulfide bonds that can be reduced by intracellular glutathione facilitating biodegradability. When glutathione was treated for longer than 12 h to the solution containing encapsulated ONCM, ONCM was detectable by ELISA (Supplementary fig. 8A, B), indicating that only free ONCM released from the nanogel complex can be measured by ELISA. We found that the amount of free ONCM in DRGs increased only slightly at 3 and 7 dpi, but very markedly at 14 dpi (Fig. 7F), demonstrating that free ONCM is slowly released from the nanogel complex enabling sustaining activity of ONCM *in vivo*. Injection of REP L-NG/ONCM did not invoke strong inflammatory responses in DRGs, and there was no evidence of significant damage to DRG neurons by the nanocomplex (Supplementary fig. 9)

### Nanogel-mediated delivery of ONCM protein to DRGs achieves robust central regeneration of dorsal column axons beyond spinal cord lesion

We then tested the effects of nanogel-ONCM complex on central axon regeneration in *in vivo* spinal cord injury model. We created dorsal hemisection at T9 spinal cord to cut ascending dorsal column sensory axons in the spinal cord (Fig. 8A). REPL-NG/ONCM(3:1) was injected into L4 and L5 DRGs 2 weeks before or 1 day after the spinal cord injury, and the extent of regeneration traced by cholera toxin B (CTB) was compared to animals with injury only, REP L-NG only, or subjected to preconditioning SNI. The dorsal hemisection resulted in retraction of CTB traced DRG sensory axons (Fig. 8B). Injection of REPL-NG alone without ONCM conjugated slightly counteracted the axonal retraction, but there were virtually no axons growing beyond the caudal injury border. In contrast, REPL-NG/ONCM injection 2 weeks before creating the spinal lesion led to a robust regeneration beyond the lesion border (Fig. 8B, C). CTB traced axons traversing the lesion site exhibited a long-range linear growth along the rostrocaudal direction reaching up to 2 mm rostral to the caudal injury border (Fig. 8B, D; Supplementary fig. 10). The mean number of regenerating axons beyond the caudal lesion border was comparable to that achieved by preconditioning SNI up to 400 ⎧m from the injury epicenter, but axons regenerating at the region more rostral to 400 ⎧m were observed only in animals with REPL-NG/ONCM injection 2 weeks before the lesion (Fig. 8C, D). To assess feasibility of the REPL-NG/ONCM as a clinically applicable therapeutic option, we injected REPL-NG/ ONCM one day after injury. Although post-injury injection also resulted in enhanced axon re generation, the extent was considerably decreased compared to the pre-injury injection (Fig. 8 B-D). Delivery of the nanocomplex to DRGs did not influence the extent of macrophage activation at the lesion site (Supplementary fig. 11). Galanin

**Figure 8.**
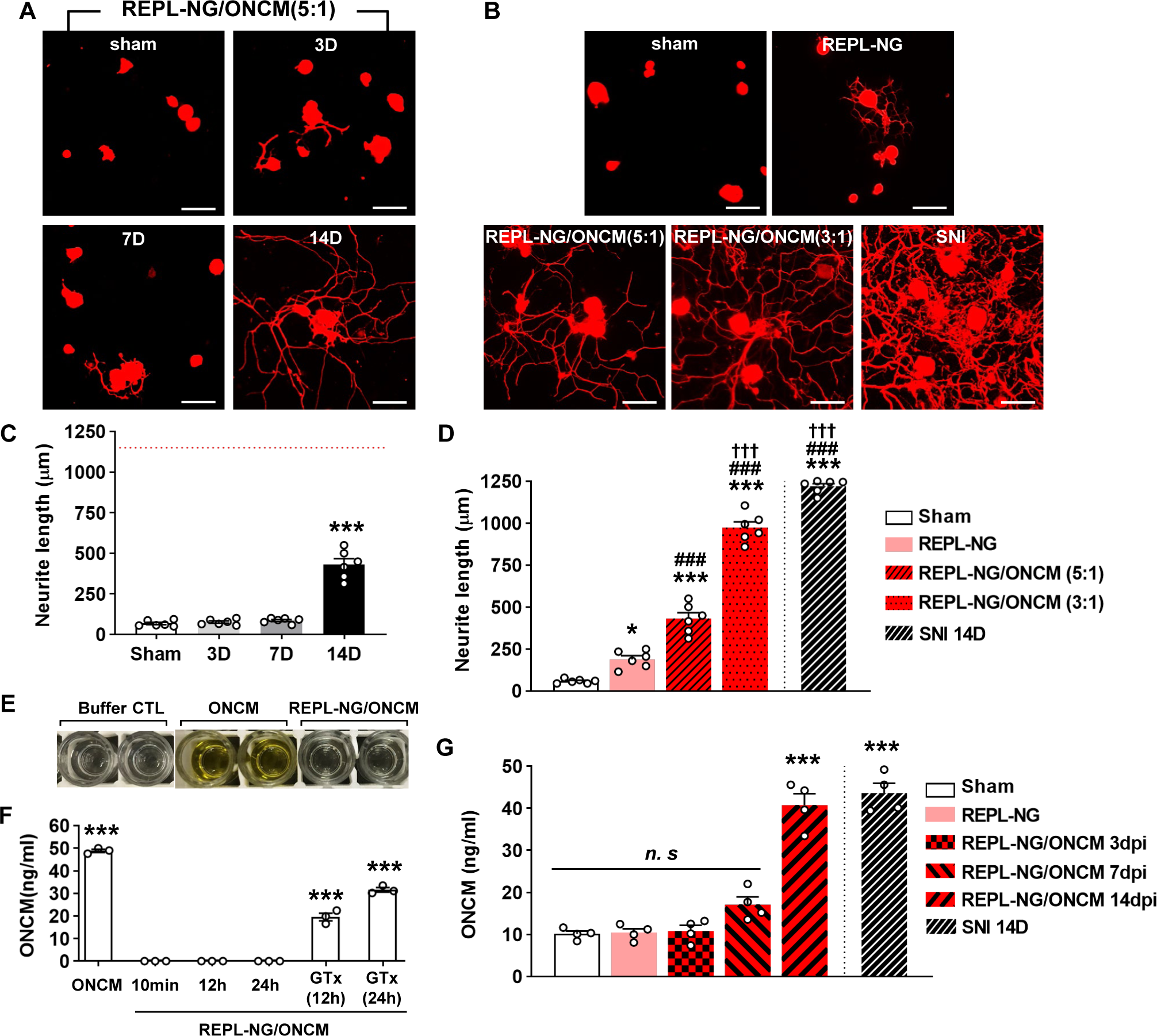
REPL-NG/ONCM complex enables sustained activity of ONCM *in vivo*. A, Representative images of neurite outgrowth of DRG neurons taken from intraganglionic in jection of REPL-NG with ONCM conjugated at 5:1 ratio, REPL-NG/ONCM (5:1) at 0 (sham), 3, 7 and 14 d after injection. Neurons from the L4 and L5 DRGs were cultured for 15 h before being fixed for the immunofluorescent visualization of neurites with anti-β III tubulin. Scale bar, 100 µm. C, Comparison of the mean neurite length between cultures. N = 6 animals p er group. \*\*\**p* < 0.001 compared with control values by one-way ANOVA followed by Tukey’s *post hoc* analysis. B, Neurite outgrowth of cultured DRGs neurons taken from intragangli onic injection of vehicle (sham), REPL-NG, REPL-NG with ONCM conjugated at 5:1 ratio; REPL-NG/ONCM (5:1) and REPL-NG with ONCM conjugated at 3:1 ratio; REPL-NG/ON M (3:1) mice at 14 d after injection. Neurons from the L4 and L5 DRGs were cultured for 15 h before being fixed for the immunofluorescent visualization of neurites with anti-β III tubulin. Scale bar, 100 µm. D, Quantification graph comparing the mean neurite length. \*\*\**p* < 0.0 01 and \**p* < 0.01 compared with sham values; ### *p* < 0.001 compared with REPL-NG value s; †††*p* < 0.001 compared with REPL-NG/ONCM (5:1) value by one-way ANOVA followed by Tukey’s *post hoc* analysis. E, Microtiter plate showing the results of ONCM-ELISA. Wel ls indicate lysis buffer (buffer CTL), lysis buffer with ONCM (ONCM), and lysis buffer with REPL-NG/ONCM (REPL-NG/ONCM) 10 min after incubation at the left of the wells. F, ELI SA measurement of oncomodulin concentration in the tissue lysis buffer treated ONCM conjugate with or without REPL-NG, recombinant ONCM, ONCM conjugate with REPL-NG, or Glutathion (GTx) at a different incubation time. N = 3 independent experiments per each group. \*\*\**p* < 0.001 compared with control values by one-way ANOVA followed by Tukey’s post hoc analysis. G, ELISA measurement of oncomodulin concentration in the DRGs obtained at 3, 7, and 14 days after intraganglionic injection of REPL-NG/ONCM. N = 4 animals per each group. \*\*\**p* < 0.001 compared with control values by one-way ANOVA followed by Tuk ey’s post hoc analysis.

**Figure 9.**
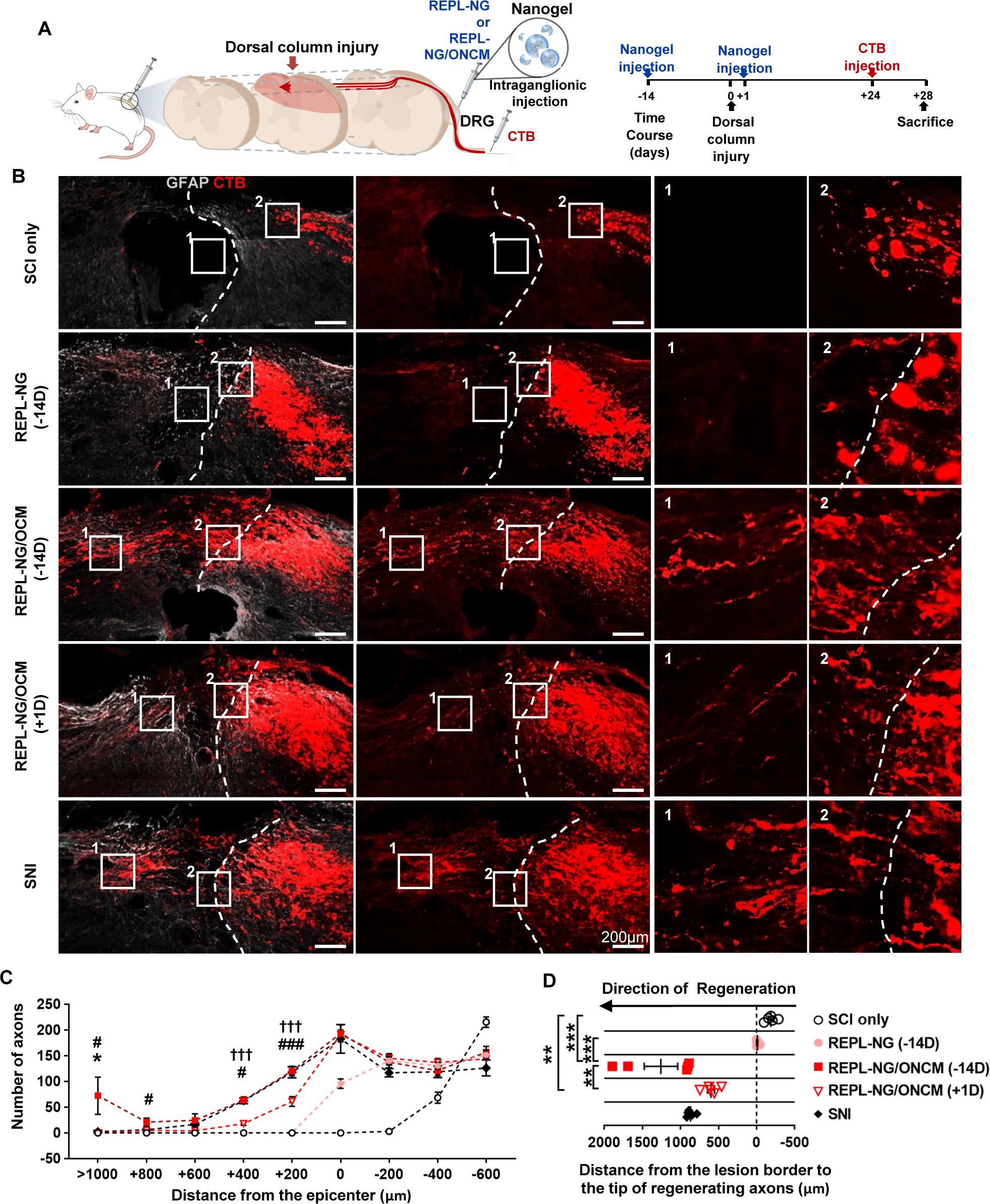
REPL-NG/ONCM complex robustly increases regeneration dorsal column axons following spinal cord injury. A, Experimental scheme depicting the time course of the experimental procedures. B, Representative images of cholera toxin B (CTB)-labeled axons (red) and GFAP-immunostained spin al cord sections (gray) from animals subjected to spinal cord dorsal hemisection only or injected with REPL-NG 14 d before injury (REPL-NG; -14D), with REPL-NG/ONCM 14 d before injury (REPL-NG/ONCM; -14 D), or with REPL-NG/ONCM 1 d after injury (REPL-NG/O NCM; +1D), and those subjected to preconditioning sciatic nerve injury (SNI) 2 weeks before creating the spinal lesion. Dashed lines indicate caudal lesion borders as determined by GF AP immunostaining. The boxed regions are magnified in the right panels. Scale bars represent 200 µm. C, Quantification graph comparing the number of CTB-positive axons within counting blocks located at different distances from the lesion border. The numbers in the x axis indicate the distance of the rostral border of a counting block from the lesion border. \**p* < 0.01 compared with SNI and REPL-NG/ONCM (-14D) values; ### *p* < 0.001 and #*p* < 0.01 comp red with REPL-NG/ONCM (-14D) and REPL-BF/ONCM (+1D) values; †††*p* < 0.001 compared with SNI and REPL-NG/ONCM (+1D) value by one-way ANOVA followed by Tukey’s *post hoc* analysis. D, Quantification graph comparing the longest distance of regenerating axons from the caudal lesion border. Individual circles plot the longest distance of each animal. Negative values indicate the degree of the retraction from the lesion border. \*\*\**p* < 0.001\*\**p*< 0.01; \**p* < 0.05; between animals (one-way ANOVA followed by Tukey’s *post hoc* analysi s). N = 6 animals for each group.

## Discussion

There is mounting evidence that non-neuronal cells in DRGs may contribute to enhanced axon growth potential by a well-characterized “preconditioning injury” paradigm (Poplawski et a l., 2018; Zigmond and Echevarria, 2019; Avraham et al., 2020; Jager et al., 2020). Among various non-neuronal cells, functional significance of macrophage involvement was thoroughly documented by either pharmacological inhibition using minocycline (Kwon et al., 2013) or genetic ablation of CCR2 (Niemi et al., 2013). Perineuronal macrophage activation in DRGs following peripheral nerve injury is also implicated in the evolution of neuropathic pain (Vega-Avelaira et al., 2009; Yu et al., 2020; Silva et al., 2021). In CNS, microglial activation surrounding axotomized neurons were linked to axonal regeneration or changes in synaptic structures (Kreutzberg, 1993; Shokouhi et al., 2010). These findings strongly suggest that perineuronal myeloid cells exert profound influence on neurons of which axon are injured at a distant site. Previous studies have identified various signals that DRG sensory neurons transmit following axonal injury to activate macrophages (Kwon et al., 2015; Simeoli et al., 2017; Bravo-Caparrós et al., 2020). However, it remains largely unknown how exactly the perineuronal macro phages affect neurons in return to support their potential of regenerating injured axons.

The present study provides several lines of evidence that ONCM is the macrophage-derived effector signal to enhance intrinsic capacity of DRG sensory neurons for axonal regeneration. First, ONCM expression was increased exclusively in DRG macrophages following S NI in a manner dependent on CCR2, a receptor for CCL2 ligand that was shown to recruit and activate macrophages (Kwon et al., 2015; Wang et al., 2018; Bravo-Caparrós et al., 2020). Second, SNI did not lead to enhancing axon growth capacity in ONCM deficient mice when DRG neurons were cultured at 7 dpi, demonstrating the necessity of ONCM. However, ONC M was not required for early induction of conditioning effects at 2 dpi, a time point when injury-induced increase in macrophages has not yet taken place (Iwai et al., 2021). This finding suggests that contribution of ONCM becomes significant only when macrophages are substantially activated. Third, ONCM deficient macrophages failed to produce MCM with neurite out growth activity in the *in vitro* neuron-macrophage interaction model. In the same model, MC M from WT macrophages co-cultured with ONCM deficient neurons contained potent neurite outgrowth activity, precluding a possibility that neuron-derived ONCM is involved in the injury-induced conditioning effects. Finally, overexpression of ONCM engineered to be released extracellularly in DRGs *in vivo* increased axon growth capacity of DRG neurons, success fully mimicking the conditioning effects. An intriguing finding was the absence of the signal sequence within the ONCM coding region, although ONCM was readily released into MCM from culture primary macrophages. It remains to be determined how ONCM is released from macrophages not utilizing the canonical secretory pathway.

We found that ONCM possesses highly robust neurite outgrowth activity *in vitro*, to the extent comparable to that achieved by preconditioning injury. When cultured DRG neuron s were exposed to other molecules such as NGF and IL-6 (used as hyper IL-6 here) that had been reported to induce neurite outgrowth (Cafferty et al., 2001; Cafferty et al., 2004), no significant enhancement was observed under the same experimental condition where ONCM potently induced neurite outgrowth. Our finding that ONCM successfully overcame inhibitory influence of CSPGs was also in contrast with the previous report showing no effect of ONCM in the *in vitro* spot gradient assay (Harel et al., 2012). These discrepancies between our and previous findings might be due to different culture conditions or a species difference between mouse (our study) and rat (the above previous studies) DRG neurons. Nonetheless, it could be argued that ONCM is the most potent molecular factor to induce neurite outgrowth in culture DRG neurons at least in our culture condition. ONCM was identified as a potent signal for axon regeneration of retinal ganglion cells (Yin et al., 2006). ONCM *in vitro* did not induce axon outgrowth when treated alone, but potently increase the axon growth when combined with mannose and forskolin, a stimulator of adenylate cyclase (Yin et al., 2006). In our study, increasing cAMP alone slightly increased the extent of neurite outgrowth. ONCM treatment alone achieved almost maximum extent of axon growth that could be expected following preconditioning injury. The precise mechanism by which ONCM potently increases neurite outgrowth in DRG neurons is presently unknown. Our RNA-seq data suggest that neuropeptide-related genes may be downstream effectors for ONCM effects on neurite outgrowth. Potential roles of the neuropeptide-related genes in sensory axon regeneration were demonstrated. For example, disruption of the *Gal* gene reduced regenerative capacity of sensory neurons (Holmes et a l., 2000). Furthermore, intrathecal administration of NPY and VIP resulted in an increase in the percentage of dissociated DRG neurons with growing neurites (White and Mansfield, 1996). Interestingly, *Gal* gene was highly expressed in retinal ganglion cells following axotomy plus lens injury in which retinal ganglion cell axon regeneration is promoted via macrophage-derived ONCM (Fischer et al., 2004). In contrast, ONCM treatment did not significantly increased the expression of hub transcription factors implicated in the condition effects. We verified persistent activation of c-Jun and c-Myc following SNI in ONCM deficient mice, indicating that ONCM is a downstream effector molecule whose expression may be influenced by those hub transcription factor following preconditioning injury.

To facilitate effective delivery of ONCM protein to DRGs, REPL-NG drug delivery system was chosen in this study. NGs are nano-sized (usually 100-200 nm) particles with physical properties of hydrogels formed by crosslinked swellable polymer networks (Sharma et al., 2016; Soni et al., 2016). Owing to its capacity to load a large amount of drug and the fluid-like deformability (Sharma et al., 2016; Neamtu et al., 2017), we speculated that NG would be an ideal carrier targeting organs of very small size like the DRG. Furthermore, NG can protect biomolecular cargos from being degradation or elimination and thereby enhance stability of target molecules exposed to biological fluid (Soni et al., 2016). After thiolation and successive oxidation of EPL, the resulting REPL-NG possesses multiple disulfide bonds and primary amines. Primary amines exhibit positive charges when REPL-NG is dissolved in a solution with pH 7.4. Ionic interaction with charged polymers and counter-charged proteins has been employed for the controlled release of protein drugs (Kang et al., 2012; Tian et al., 2013). The positive charge in REPL-NG is exploited to form an electrostatic bond with negatively charged ONCM with pI value 3.9. This electrostatic force will allow nano-sized complexation between REPL-NG and ONCM and would imparts great stability of ONCM *in vitro* with serum com ponents and *in vivo* DRG tissue. Indeed, this REPL-NG/ONCM complex completely hindered antibody binding to ONCM even in the presence of detergent-containing lysis buffer, resulting in no detection of ONCM in ELISA assay. Strong and stable encapsulation of ONCM was also supported by the observation that free ONCM level in DRGs was slowly increased by 1 4 days after being injected as REPL-NG/ONCM complex. REPL-NG/ONCM is supposed to be delivered within cells of interest as a form nanocomplex. Glutathione-triggered reduction of disulfide bonds is often utilized for intracellular drug delivery in a controlled manner (Chi et al., 2017; Feng et al., 2018; Qiu et al., 2020). Multiple disulfide bonds in REPL-NG can be degraded by intracellular glutathione releasing free ONCM within cytoplasmic compartment. Our finding that addition of glutathione burst REPL-NG/ONCM enabling detection of ONC M by ELISA supports the notion that intracellular glutathione would be a key mechanism for efficient intracellular delivery of ONCM. Collectively, our study demonstrated that encapsulation of ONCM by REPL-NG could greatly improve stability and prolong its bioactivity in DR G tissue.

We observed that NG-mediated delivery of ONCM promoted regeneration of sensory axons well beyond spinal cord lesion, to the extent surpassing that observed following preconditioning injury. The longest distance of regenerating axons from the lesion center was almost 2 mm, and this amount of regeneration would be certainly notable when compared to our ow n experiences with overexpression of a chemokine or a transcription factor in the exact same injury model (Kwon et al., 2015; Shin et al., 2020). We speculate that this remarkable level of axon regeneration was due to enhanced stability and sustained bioactivity of ONCM protein in DRG tissue. As exemplified by inhibition of ONCM activity by serum components in this study, peptide drugs like growth factors known to stimulate axon regeneration would be susceptible to various molecular factors within tissue environment and it would be challenging to maintain their bioactivity in a prolonged time scale. The stability issue would be even more pivotal in axon regeneration studies since cellular processes of axon regeneration require quite a long time-scale with at least a couple of weeks in rodents. One caveat is that stability of a peptide drug should be optimized for the best outcome. In this study, we found that injection of REPL-NG/ONCM one day after spinal cord lesion did not lead to axon regeneration as robust as the same injection 2 weeks before the lesion. Based on the temporal course of ONCM level in DRGs following injection, it is conceivable that the availability of free ONCM may not be sufficient for the duration of initial one or two weeks following spinal cord lesion, when key biological mechanisms for axon regeneration should be fully mobilized to stimulate elongation of cytoskeletal proteins. Further studies will address this optimization issue to produce the best outcome by post-injury injection of REPL-NG/ONCM within a clinically relevant therapeutic window.

## Materials and methods

### Animals

Adult female Sprague Dawley rats (250–300 g) and C57BL/6 wild-type (WT) mice were purchased from Orient Bio Inc. ONCM- and CCR2-deficient mice on C57BL/6 background were purchased from European Mouse Mutant cell repository (MGI ID 4431716) and The Jackson Laboratory (stock 004434), respectively. Cx3cr1-GFP mice on C57BL/6 background we re purchased from Jackson Laboratory (stock 005582). The Institutional Animal Care and Us e Committee of Ajou University School of Medicine approved all animal protocols.

### Surgical procedures

Mice were anesthetized with intraperitoneal injection of the ketamine and xylazine mixture (1 00mg/kg and 10mg/kg, respectively). For creation of SNI, muscles were displaced to expose t he right sciatic nerve, and the nerve was ligated proximal to its trifurcation. The sciatic nerve was completely transected below the ligation site with fine surgical scissors. Rats were anesthetized with intraperitoneal injection of the ketamine and xylazine mixture (90mg/kg and 8mg/ kg, respectively). SNI in rat was performed following the same procedures as for mice. To create a dorsal column lesion in the spinal cord, a dorsal laminectomy was performed at the T9 level and bilateral dorsal columns with adjacent lateral columns were cut out with iridectomy scissors. To visualize regenerating axons, cholera toxin subunit B (CTB; List Biological Laboratories) was injected using a protocol modified slightly from that in the previous report (Kw on et al., 2015). Briefly, after the sciatic nerve between the thigh muscles was exposed, a small incision was made on the perineurium just proximal to the trifurcation site. Two microliters of unconjugated CTB solution (1% in PBS) were slowly injected using the Hamilton syringe through the perineural incision and animals were killed 5 d after the injection.

### Primary culture of dissociated adult DRG neurons and neurite outgrowth assay

DRGs were freshly dissected and treated with 125 U/ml type XI collagenase (Sigma-Aldrich) dissolved in DMEM (Hyclone) for 90 min at 37°C with a gentle rotation. After washing five times with DMEM, cells were dissociated by trituration using a pipette tip and centrifuged at 1500 rpm for 3 min. Cell pellets were resuspended in Neurobasal-A (Invitrogen) supplemented with B-27 (Invitrogen) and plated onto eight-well culture slides (BD Biosciences) precoated with 0.01% poly-D-lysine (Sigma-Aldrich). The culture slides were incubated with 3 ⎧g/ml aminin (Invitrogen) for 2 hours at 37°C before cell plating. When assessing neurite outgrowth under growth-inhibitory environment, the culture slides were coated with 1.0 ⎧g/ml chondroi tin sulfate proteoglycans (CSPGs; Millipore) together with laminin. The culture duration was strictly limited to 15 h, at which time point only negligible neurite outgrowth is observed in control condition (without conditioning injury). For treatment of NGF and hyper IL-6, cells we re treated with NGF (Life technologies) or hyper IL-6 (R&D systems) for 24 h immediately after cell plating. For co-treatment of NGF and hyper IL-6, cells were treated with NGF (Life technologies) for 6 h, then treated with hyper IL-6 (R&D system) for 18 h. For treatment of Forskolin (15⎧M; Sigma-Aldrich), D-mannose (1 M; Sigma-Aldrich), dibutyryl-cAMP (db-cAMP; 100⎧M; Calbiochem) and 100 ⎧g/ml of recombinant ONCM, cells were treated with forskolin, mannose and recombinant ONCM or db-cAMP and recombinant ONCM for 24 h immediately after cell plating. Neurite outgrowth was visualized by immunostaining with mouse anti-β III tubulin (1:1000; Promega) followed by incubation with Alexa 594-conjugated anti-mouse secondary antibodies (Invitrogen).

### Primary bone marrow-derived macrophage culture

To harvest bone marrow-derived macrophages (BMDMs), adult mice were anesthetized with an overdose of ketamine and xylazine mixture. Bilateral femurs and tibias were flushed using 26-gauge needles into DMEM containing 10% FBS (Hyclone). Bone marrow cells were collected and centrifuged at 1500 rpm for 5 min. Cell pellets were resuspended in red blood cell lysis buffer (0.15 mol/l NH_4_Cl, 10 mmol/l KHCO_3_, and 0.1 mmol/l Na_2_EDTA, pH 7.4). Purified bone marrow cells were centrifuged at 1500 rpm for 3 min and filtered through a 100 μm cell strainer (BD Biosciences). The bone marrow cells were plated onto a 100 mm culture dis h or six-well plate (BD Biosciences) coated with 0.01% poly-D-lysine and cultured in DME M supplemented with 1% penicillin/streptomycin and 10% FBS. Macrophage colony-stimulating factor (M-CSF; Invitrogen) was added at a concentration of 20 ⎧g/ml to facilitate differentiation of the bone marrow cells into macrophages (7–10 days).

### Neuron-macrophage co-culture

Neuron-macrophage co-cultures were established following the methods described previously (Yun et al., 2018) with slight modifications. Dissociated DRG neurons were incubated in a 6-well plate with DMEM supplemented with B-27 for 4 h. Then, BMDMs were plated on the transwell inserts placed on top of the 6-well plate wells at a ratio of 1:5 (neurons to macrophages). Four hours after setting up the co-cultures, cells were treated with dibutyryl-cA MP (db-cAMP; 100μM; Calbiochem) or PBS as a control. After 24 h, BMDMs grown on the cell culture inserts were transferred to a new culture plate. The culture medium was replaced with fresh DMEM supplemented with B-27. The co-cultures were maintained for 72 h without changing the medium and then MCM was collected, centrifuged at 1500 rpm for 5 min, and passed through a 0.2 μm filter (BD Biosciences) to remove any remaining cellular debris. For neurite outgrowth assays with MCM, primary dissociated DRG neurons were plated on eight-well culture slides as described above and maintained in culture for 2 h to allow for attachment to the culture dishes before replacing the medium with the collected MCM. After 15 h in culture, the cells were fixed and immunostained for β III tubulin to visualize neurite outgrowth.

### Quantitative measurement of mRNA by real-time PCR

Total RNAs were extracted from dissected DRGs or cultured cells using Trizol (Invitrogen) according to the manufacturer’s protocol. The amount of RNA was determined using NanoDrop Lite Spectrophotometer (Thermo Fisher Scientific) at 260 nm. Two microgram of RNA was reverse-transcribed to cDNA using a standard RT protocol. One microgram of cDNA was added to the PCR-reaction premix (Takara Bio) with 10 pM corresponding primer pairs. The following primers were used for PCR: 18S ribosomal RNA (rRNA), 5’-CGGCTACCACATCC AAGGAA-3’ (forward), 5’-TGCTGGCACCAGACTTGCCCTC-3’ (reverse); ONCM, 5’-TT CTGAGCGCTGATGACATT-3’ (forward), 5’-CGCTCTGGAACCTCTGTAGG-3’ (reverse); NPY, 5’-TGGACTGACCCTCGCTCTAT-3’ (forward), 5’-TGTCTCAGGGCTGGATCTC T-3’ (reverse); Galanin, 5’-GTGACCCTGTCAGCCACTCT-3’ (forward), 5’-GGTCTCCTT TCCTCCACCTC-3’ (reverse); Sprr1a, 5’-CCCCTCAACTGTCACTCCAT-3’ (forward), 5’-CAGGAGCCCTTGAAGATGAG-3’ (reverse); Vip, 5’-TTCACCAGCGATTACAGCAG-3’ (forward), 5’-TCACAGCCATTTGCTTTCTG-3’ (reverse); Map2, 5’-CTGGACATCAGCC TCACTCA-3’ (forward), 5’-AATAGGTGCCCTGTGACCTG-3’ (reverse); CD68, 5’-ACTC ATAACCCTGCCACCAC-3’ (forward), 5’-.GATTTGAATTTGGGCTTGGA-3’ (reverse); Glutamine synthetase, 5’-GGGGTGATAGCAACCTTTGA-3’ (forward), 5’-ACTGGTGCC TCTTGCTCAGT-3’ (reverse); GFAP, 5’-GCTTCCTGGAACAGCAAAAC-3’ (forward), 5’-AAGGTTGTCCCTCTCCACCT-3’ (reverse); P0, 5’-GGTTTACACGGACAGGGAAA-3’ (forward), 5’-GTCCCTTGGCATAGTGGAAA-3’ (reverse); S100, 5’-CCATGGAGACC CTCATCAAT-3’ (forward), 5’-TTGAAGTCCACTTCCCCATC-3’ (reverse). Quantitative R T-PCR was performed according to the protocol supplied by the SYBR Green PCR kit using the 7500 Real-Time PCR System (Applied Biosystems). Cycling conditions were 94°C for 30 s, 55°C ∼ 64°C for 30 s, and 72°C for 60 s, with a total of 40 cycles. Melting curves were generated after the last extension step, and the CT values were quantified by the Applied Biosystems 7500 software. Target gene expression was normalized with the expression of 18S rRN A as an internal control.

### ELISA

To quantify the amount of ONCM in DRG tissue, L4 and L5 DRGs were dissected and lysed to extract proteins, and the ONCM concentration was measured using an ELISA kit for oncomodulin (catalog #MBS908312; Mybiosource), according to the manufacturer’s protocol. In each well, 50 μl of DRG lysate at a concentration of 100 ng/μl was dispensed. Each experiment was performed in triplicate. To measure the concentration of ONCM in MCM obtained from cultured BMDMs, MCM was centrifuged at 1500 rpm for 5 min and passed through a 0.2 μm filter (BD Biosciences) to remove remaining cellular debris. One hundred microliter of C M was dispensed in each well. Three independent cultures were performed to collect CM for each condition, and each CM was measured in triplicate. To measure the protein level of ON CM in protein lysis buffer treated ONCM conjugate with or without REPL-NG, recombinant ONCM, ONCM conjugate with REPL-NG, or Glutathion (10 mM; Sigma-Aldrich) treated with RIPA Lysis buffer (Thermo Fisher) for 10 min, 12 h, or 24h at room temperature. 50 micro liter of each group was dispensed in each well. Four independent treatment experiments were performed for each condition, and each treatment condition was measured in duplicate.

### ONCM expression vector and transfection

pCMV-ONCM was constructed by cloning ONCM cDNA as BamHI/NotI fragment from My c-DDK-tagged oncomodulin (OriGene) in a mammalian expression vector pCMV (Addgene). To prepare a construct with the signal sequence (SS) for secretion of ONCM by translocation across the endoplasmic reticulum membrane, the N-terminal bacterial alkaline phosphatase (PhoA) signal peptide sequences (MKQSTIALALLPLLFTPVTKAR) were subcloned from pI g20 vector. PhoA signal peptide was amplified using a 5’-ATGAAGCAGTCCACCATCGC PCR primer and 5’-GCGGGCCTTGGTCACGGGGG PCR primer, and the PCR product was subcloned into Strataclone blunt PCR vectors (Stratagene) and subcloned further into pCM V-ONCM to generate the pCMV-SS-ONCM construct. PC12 cells were cultured in DMEM containing 10% horse serum and 1% penicillin-streptomycin. PC12 cells (1.5 · 10^5^ cells) were seeded on PDL-coated 6 well plates and incubated for 24 h. To induce neuronal differentiation, the medium was changed to DMEM containing 1% horse serum, 1% penicillin-streptomycin, and 100 ⎧g/ml NGF. The cells were further cultured for 6 d, and the differentiation medium was replaced every 2 d. HEK293T cells and Raw 264.7 cells were maintained in DMEM supplemented with 10% fetal bovine serum (FBS) and 1% penicillin-streptomycin. Cells were transfected with Lipofectamine 2000 (Thermofisher) following the manufacturer’s instructions. Briefly, cells were transfected at 60–80% confluency in 6-well plates using 2 μg of the expression plasmid. After 48 hr, cells were washed with PBS and lysed by applying 250 μl lysis buffer (20 mM HEPES at pH 7.5, 100 mM KCl, 5 mM MgCl_2_, 1 mM dithiothreitol, 5% glycerol, and 0.1% Triton X-100 supplemented with Roche Protease Inhibitor cocktail) and then roc ked for 10 min at 4°C. The resulting cell lysate was divided into aliquots for further analysis.

### Preparation and injection of recombinant adeno-associated virus

Recombinant adeno-associated virus (AAV) vector containing a CCL2 expression cassette w as generated by replacing GFP cassette of pAAV-CAG-GFP (Addgene) with the full-length c DNA for CCL2 (OriGene). AAV serotype 5 (AAV5) virus preparation was performed at the University of North Carolina at Chapel Hill Vector Core facility. Control AAV5-GFP was purchased from the same facility. The titer of both viruses was ∼1 x 10^12^ genome copies/ml. For intraganglionic injection, the L5 DRG was exposed after removal of the lateral process of the L5 vertebral bone. A Hamilton syringe configured with a glass pipette was controlled using a micromanipulator, and the tip of the glass pipette was slowly advanced into the L5 DRG under a surgical microscope; 2 μl of AAV5-CCL2 or AAV5-GFP was injected into the L5 DRG at a rate of 0.25 μl/min using a nanoinjector (#M3301R, World Precision Instruments). The recombinant AAV vector containing ONCM or SS-ONCM expression cassette was generated from pCMV-ONCM and pCMV-SS-ONCM. AAV5 viral particles containing either ONCM o r LS-ONCM were prepared at the Korea Institute of Science and Technology virus facility. T he titer of both viral particles was ∼1 x 10^14^ genome copies/ml. Intraganglionic injection was done using the same procedures described above.

### Isolation and sequencing of RNA from cultured DRG neurons

Cultured DRGs neurons were established following the methods described previously with slight modifications. For further purification, cell suspension was layered on a BSA cushion (10% w/v in Neurobasal-A) and centrifuged at 600 g for 13 min at a low brake. Myelin debris and lighter non-neuronal cells were settled at the top layer of 10% BSA, whereas the neurons were pelleted down. Cell pellets were resuspended in Neurobasal-A (Invitrogen) supplemented with B-27 (Invitrogen) and plated onto a six-well plate. Immediately after setting up the cultured DRG neurons, cells were treated with 100 ⎧g/ml of recombinant ONCM. RNA was extracted using RNeasy Plus Micro Kit (Qiagen) following the manufacturer’s protocol. Purified RNA samples at a concentration of 5200-500 ng/μL for 20 μL and RNA Integrity Number (RIN) above 9.5 were processed for library preparation and RNA-sequencing at the Macrogen. cDNA libraries were prepared using Trueseq Stranded mRNA Prep Kit (Illumina). The cDN A libraries were then sequenced using the Illumina Novaseq 6000 platform with 150 bp paired end reads. Gene ontology (GO) analysis for differentially expressed genes (n=167, PBS vs. ONCM, fold difference on log2 scale > 2) was performed by using DAVID software (https://david.ncifcrf.gov). For neuropeptide-related signatures in the GO database, gene set enrichment analysis (GSEA) was performed using GSEA (https://www.gsea-msigdb.org/gsea).

### Synthesis and characterization of reducible ε−poly(l-lysine) nanogel (REPL-NG)

REPL-NG was synthesized by a two-step chemical reaction: the thiolation of ε-poly(_L_-lysine) (EPL, Zhengzhou Bainafo Bioengineering, China) by 2-iminothiolane (IT, Sigma. USA) and then formation of multiple disulfide bonds among thiols in thiolated EPL. In detail, after EP L (200 mg, 42.6 μmol) and IT (213 μmol; five equivalents to the moles of EPL) were dissolved in 8 mL and 1 mL of Dulbecco’s phosphate buffered saline (DPBS; pH 7.4), respectively, two reactant solutions were mixed and then reacted to thiolate some primary amines of EPL f or 24 h at room temperature (RT). Then, 3 mL of DMSO was added into the thiolated EPL solution to form disulfide bonds, and the reaction solution was stirred for additional 24 h. After transferring the resulting REPL-NG-containing solution into a dialysis membrane (molecular weight cut-off 25 kDa), the REPL-NG was purified by a dialysis against deionized water for 48 h. The dialysate was filtrated to remove unwanted aggregates by a paper filter and was lyophilized. In the resulting REPL-NG, the number of the IT-derived groups was approximately 5.4 per a single EPL polymer chain and was detected by 500 MHz Bruker ^1^H-NMR spectrometer (Bruker, USA). The particle size and zeta-potential of REPL-NG were 11.0 ± 1.4 nm (polydispersity index = 0.224) and 43.9 ± 6.1 mV, respectively, and were monitored by a zeta-sizer (ELS-Z, Photal Otsuka Electronics Co., Japan). The synthesized REPL-NG was stored as a dried solid at -20 °C prior to use.

### Generation of REPL-NG/ONCM complex and intraganglionic injection of the nanocomplex

Five μg of REPL-NG (5 mg/ml in PBS solution) was mixed with 1.7 μg of recombinant ONC 1. M. To increase electrostatic interaction between recombinant ONCM and REPL-NG, REPL-NG/ONCM complex was subjected to vertical tapping at a weight ratio of 1:3 (REPL-NG to recombinant ONCM). For stabilization of REPL-NG/ONCM complex formation, REPL-NG and ONCM mixture were incubated for 30 min at RT. Immediately after the stabilization of Rl of REPL-NG/ONCM complex or REPL-NG was injected into the L4 and L5 DRG at a rate of 0.25 μl/min using a nanoinjector (#M3301R, World Precision Instruments). For intraganglionic injection, the L4 and L5 DRGs were exposed after removal of the lateral process of the L4-5 vertebral bone. A Hamilton syringe configured with a glass pipette of which tip diameter was less than 70 μm was slowly advanced into the L4 and L5 DRGs sequentially under a surgical microscope. After injection, the syringe was kept in situ for 1 min to prevent regurgitation of the injected REPL-NG or REPL/NG-ONCM complex through the injection site.

### Tissue processing and immunohistochemistry

Rats or mice were anesthetized with ketamine/xylazine mixture and perfused with heparinized PBS followed by 4% PFA in 0.2 M PB. DRGs or spinal cord tissues containing the lesion site were dissected and postfixed in 4% PFA for 2 h, followed by cryoprotection in a graded series of sucrose solutions. DRGs were cryosectioned at 12 μm thickness. For spinal cord tissue, parasagittal cryosections (at 40 μm thickness) were made in a 1:5 series. Tissue sections were mounted onto Super Frost plus slides (Thermo Fisher Scientific) and stored at -20°C until u se. DRG sections underwent antigen retrieval with 0.1 M EDTA Tris buffer, pH 9.0, at 98°C and were treated with 10% normal goat serum and 0.3% Triton X-100 for 1 h, and then the primary antibodies, dissolved in the same blocking solution, were applied at 4°C overnight. The primary antibodies were rabbit anti-GFAP (1:500; Dako), chicken anti-GFP (1:1000; Abcam), goat anti-CTB antibodies (1:10000; List Biological Laboratories), rabbit anti-Iba-1 (1:500; Dako), rabbit anti-galanin (1:1000, Millipore), rabbit anti-c-Myc (1:250, Abcam), rabbit anti-Sprr1a (1:1000; Abcam), rabbit anti-phospho-c-Jun (1:100; Millipore). Tissue sections were washed thoroughly and then incubated with appropriate secondary antibodies tagged with AlexaFluor-488 or -594 (1:500; Invitrogen) for 1 h at room temperature. The coverslips were mounted onto slides with glycerol-based mounting medium (Biomeda). Tissue sections were incubated for 30 min with NeuroTrace 640/660 for Deep-Red Fluorescent Nissl stain (1:300; Invitrogen), and underwent three wash steps of 5 min each in PBS, followed by mounting. The images were taken using an LSM-800 confocal microscope (Zeiss) and Axio scan Z1 (Zeiss).

### Quantitative image analysis

To quantify intensity of IBA-1 immunoreactivities, two ROIs each covering 150 mm^2^. All images were obtained using the same detector setting. After images were adjusted using the pre determined threshold setting using ImageJ software (available at: http://imagej.nih.gov/ij/), in tegrated intensity of the immunoreactivity per unit area was obtained. For neurite outgrowth assays, the mean neurite length per neuron was measured to compare the extent of neurite outgrowth between different experimental conditions. Neurite length was measured using Metam orph Imaging Software. Each well was divided into four quadrants, and a 200· magnification image was obtained at the center of each quadrant (four images in each well). Measurements were made in exported eight-bit TIFF files and neurite length was determined as follows: the MetaMorph “draw” function was used to draw a line with neurite and the values were calibrated in micrometers using the MetaMorph “calibrate” function. To quantify the extent of dorsal column axon growth after injury, two consecutive parasagittal sections collected at a distance of 200 ⎧m from the intersection with the caudal lesion border and containing CTB-traced axons were used for analysis. After the caudal lesion border was identified using GFAP staining, lines perpendicular to the longitudinal axis were drawn at 200 µm intervals from the lower lesion border to delineate counting blocks, with blocks named according to their shortest distance to the lesion border. The number of CTB positive axons between successive lines was counted and recorded as the number of axons in the block. The values were averaged from the sections with visible axons in each animal. In addition, the longest distance of regenerating axons beyond the caudal lesion border was recorded for each animal.

### Fluorescence-Activating Cell Sorting (FACS) for isolation of macrophages

Biolateral SNI was performed in Cx3cr1-GFP mouse. Seven days after injury, L3, 4, and 5 D RGs were freshly dissected and treated with 125 U/ml type XI collagenase (Sigma-Aldrich) dissolved in DMEM (Hyclone) for 90 min at 37°C with a gentle rotation. After washing five times with DMEM, cells were dissociated by trituration using a pipette tip and centrifuged at 1 500 rpm for 3 min. Cell pellets were resuspended in Neurobasal-A (Invitrogen) supplemented with B-27 (Invitrogen). For further purification, Cell suspension was layered on a 30% BSA and centrifuged at 600 g for 10 min at a low brake and repeat BSA cushion methods 2 more times with 15% BSA and 10% BSA, respectively. Cell pellets were resuspended in HBSS (Hyclone). The GFP positive and negative fractions were collected separately in 15 ml comical tube by fluorescence-activated cell sorter (FACS) analysis with a FACSAria III (Becton Dickinson). Each tube was centrifuged at 1500rpm for 10min at 4 °C. Total RNA was extracted from positive and negative fraction cells using the methods describe above.

### Statistical analysis

All numerical values and error bars in the quantification graphs are expressed as mean SE1. M. Statistical comparison of mean values was performed using unpaired Student’s t tests or one-way ANOVA followed by Tukey’s *post hoc* tests. Quantification graphs were generated using GraphPad Prism version 8.00 (GraphPad Software).

### ONCM

NGF + Hyper IL-6 ONCM versus SNI

ONCM + Forskolin + mannose ONCM only

cAMP only

---

**Supplementary figure 1.**
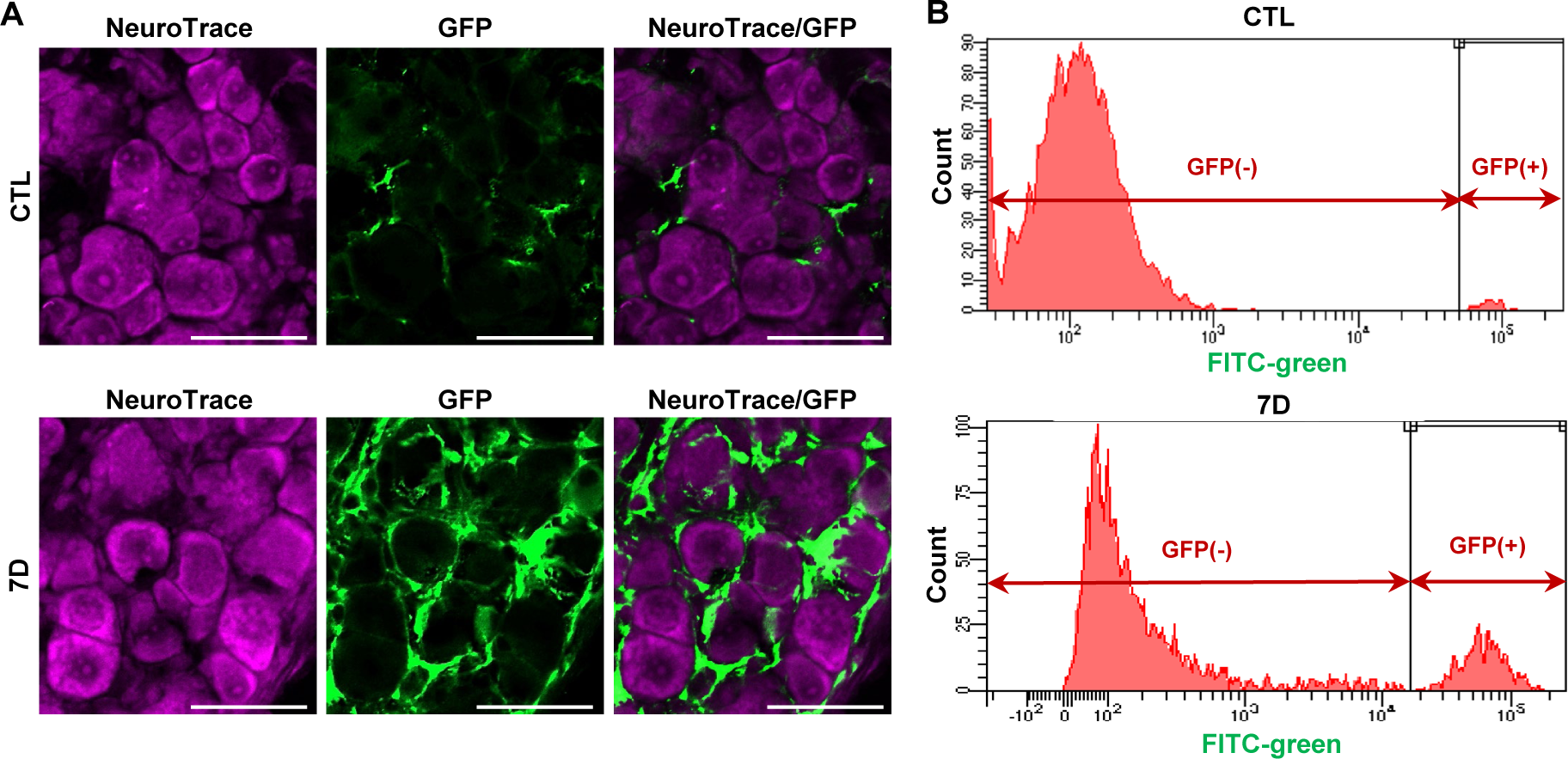
Colocalization of Cx3cr1-GFP with IBA1 and flow cytometric analysis. A, Confocal images of L5 DRG sections doubly stained for GFP (green) and NeuroTrace (vio let) at 0 (CTL) and 7 d (7D) after sciatic nerve injury. Scale bars represent 50 µm. B, Representative histogram of the flow cytometry analysis of the GFP fluorescence in dissociated DRG cells from Cx3cr1-GFP mice following sham injury (CTL) and 7 d after injury (7D).

**Supplementary figure 2.**
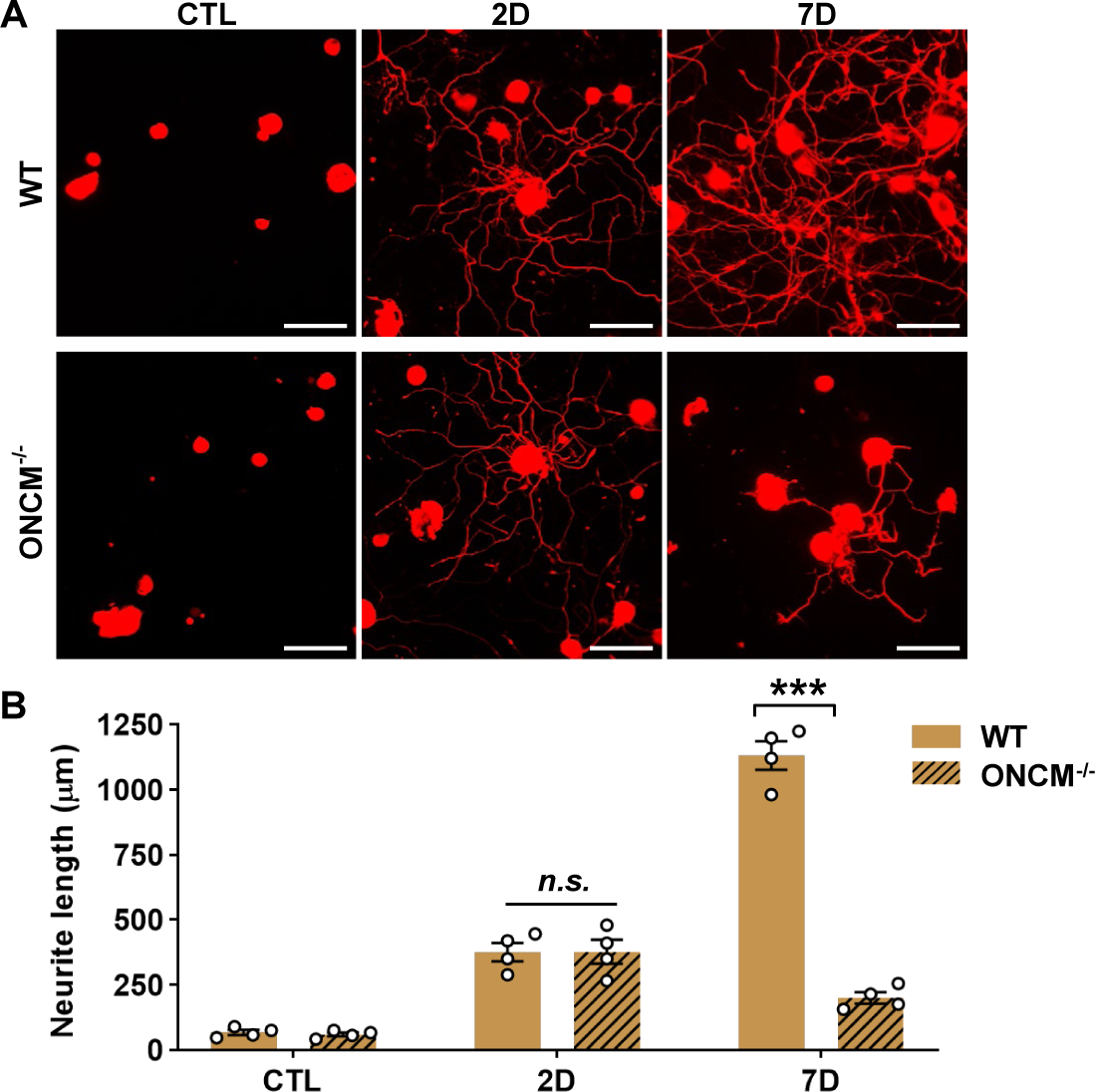
Preconditioning effects are not affected in ONCM-deficient mice at an earlier time point. A, Representative images of neurite outgrowth of DRG neurons taken from WT and ONCM^-/-^ mice at different time points after sciatic nerve injury. DRG neurons from the L3, L 4, and L5 DRGs were dissociated and cultured for 15 h before being fixed for the immunofluorescent visualization of neurites with anti-β III tubulin. Scale bars represent 100 μm. B, Comparison of the mean neurite length between cultures from WT and ONCM^-/-^ mice 0 (CTL), 2 (2D), and 7 d (7D) after injury. N *=* 4 animals for each condition. \*\*\**p* < 0.001 by unpaired *t* test.

**Supplementary figure 3.**
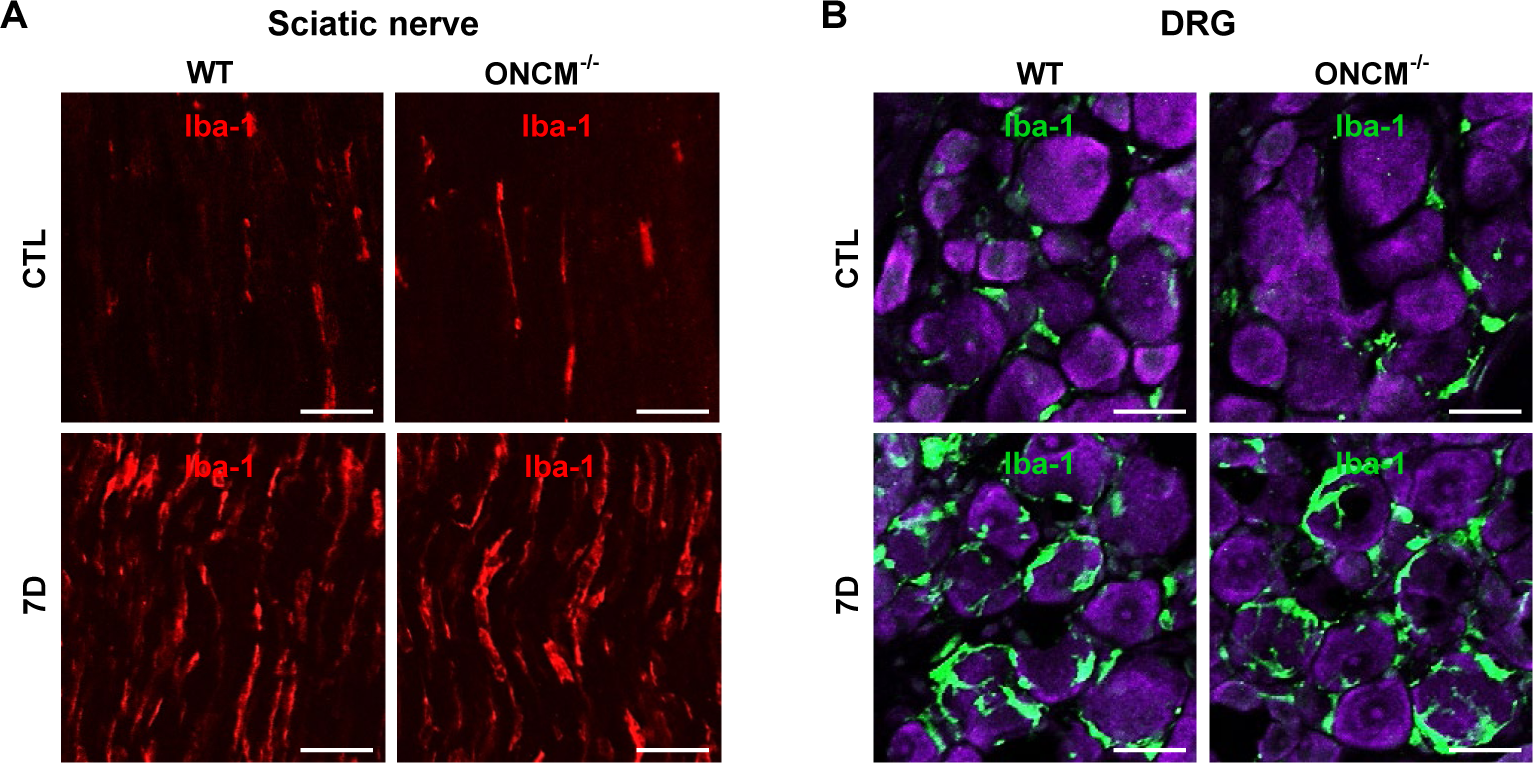
Comparison of macrophage activation at the injury site and D RG between WT and ONCM deficiency. A, Representative images of Iba-1-immunostained sciatic nerve sections obtained from animals subjected to 0 (CTL) or 7 d after sciatic nerve injury. Scale bars represent 50 μm. B. Confocal images of the L5 DRG sections obtained from WT and ONCM^-/-^ mice doubly stained for Iba-1 (green) and NeuroTrace (violet) sham injury (CTL) and 7 d after injury (7D). Scale bars represent 50 µm.

**Supplementary figure 4.**
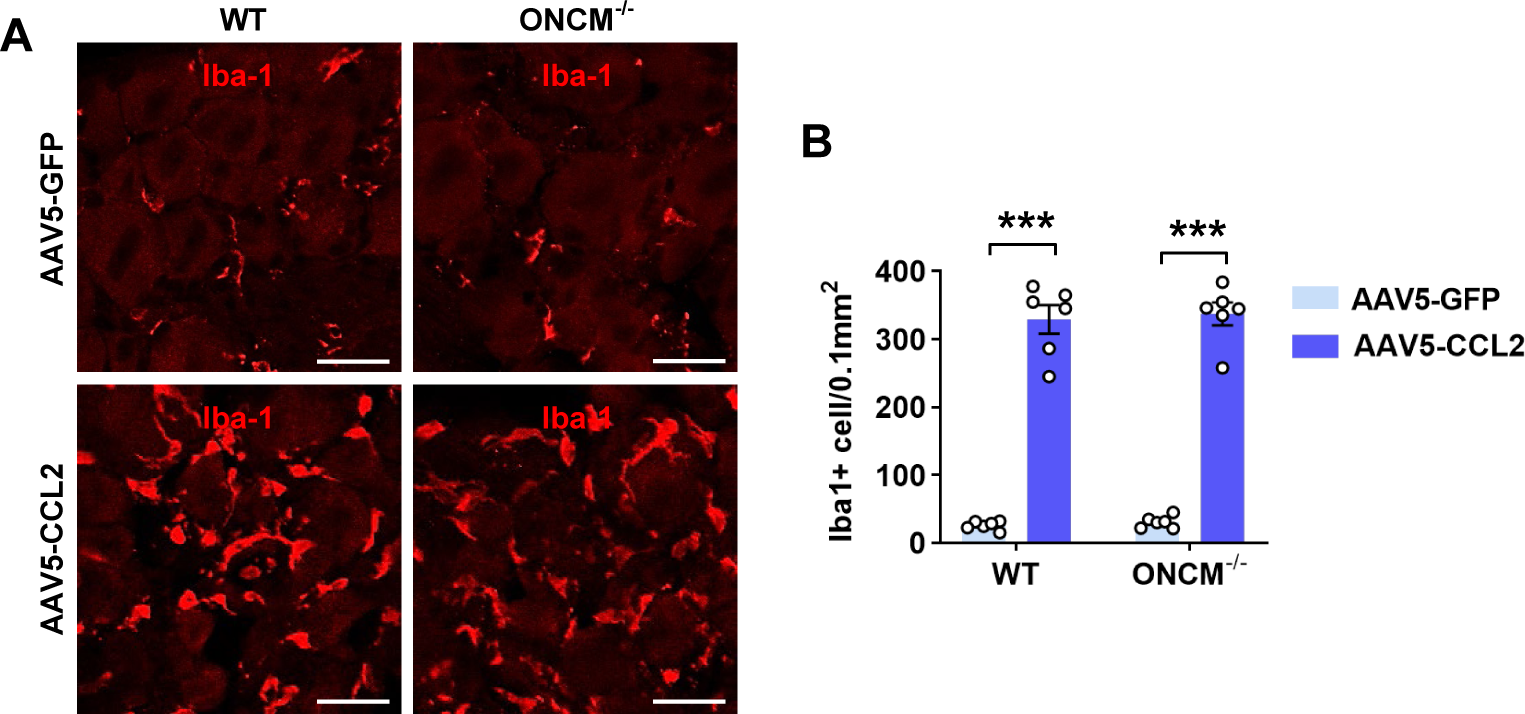
Intraganglionic AAV5-CCL2 injection increases the number of macrophages irrespective of the genotype. A, Representative images of Iba-1 staining in L5 DRG sections obtained from WT and ONC M^-/-^ mice at 28 d after intraganglionic injection of AAV5-GFP or AAV5-CCL2. Scale bars re present 50 µm. B, Comparison of the number of macrophages. N = 6 animals per group. \*\*\**p*< 0.001 between AAV5-GFP and AAV5-CCL2 injection groups by one-way ANOVA followed by Tukey’s *post hoc* analysis.

**Supplementary figure 5.**
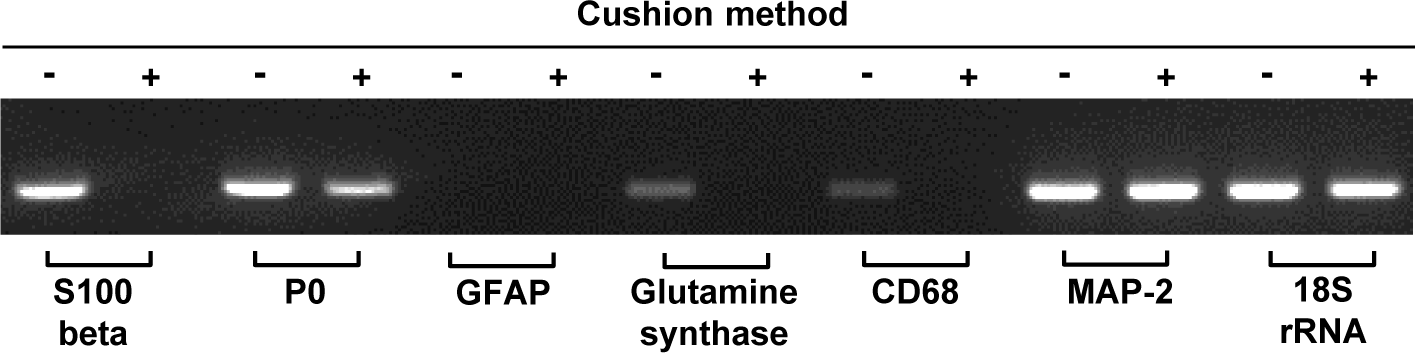
BSA cushion diminished expression of non-neural cell-specific genes. Representative images of electrophoresed RT-PCR products of various gene expression from DRG cell fractions purified by with (+) or without (-) BSA cushion method. 18S rRNA was used as an internal reference.

**Supplementary figure 6.**
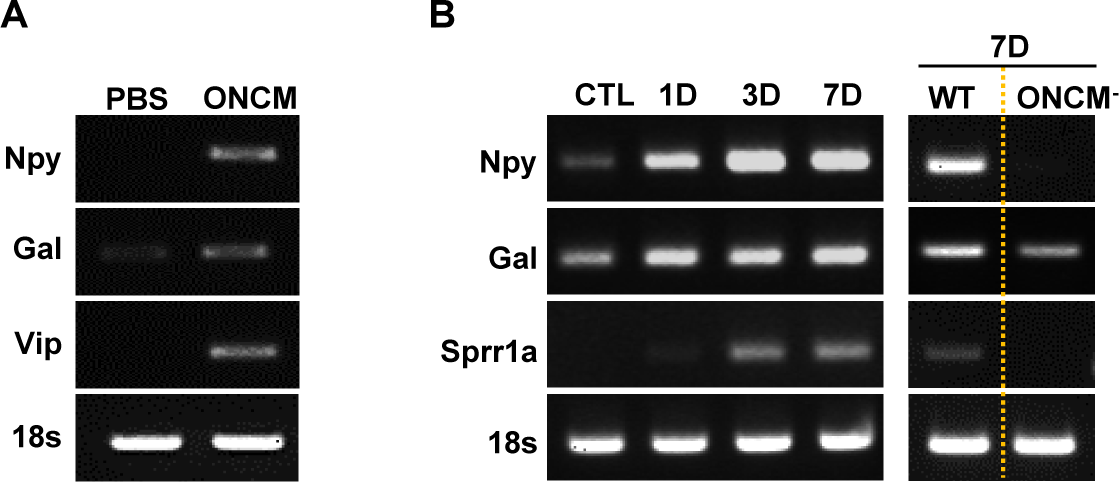
Validation of the neuropeptide gene upregulation in DRG neurons by ONCM. A, Representative images of electrophoresed RT-PCR products from cultured DRGs treated with PBS or ONCM. 18S rRNA was used as an internal reference. B, Representative images of electrophoresed RT-PCR products. 18S rRNA was used as an internal reference. DRG samples were obtained 0 (CTL), 1, 3, and 7 d after sciatic nerve injury.

**Supplementary figure 7.**
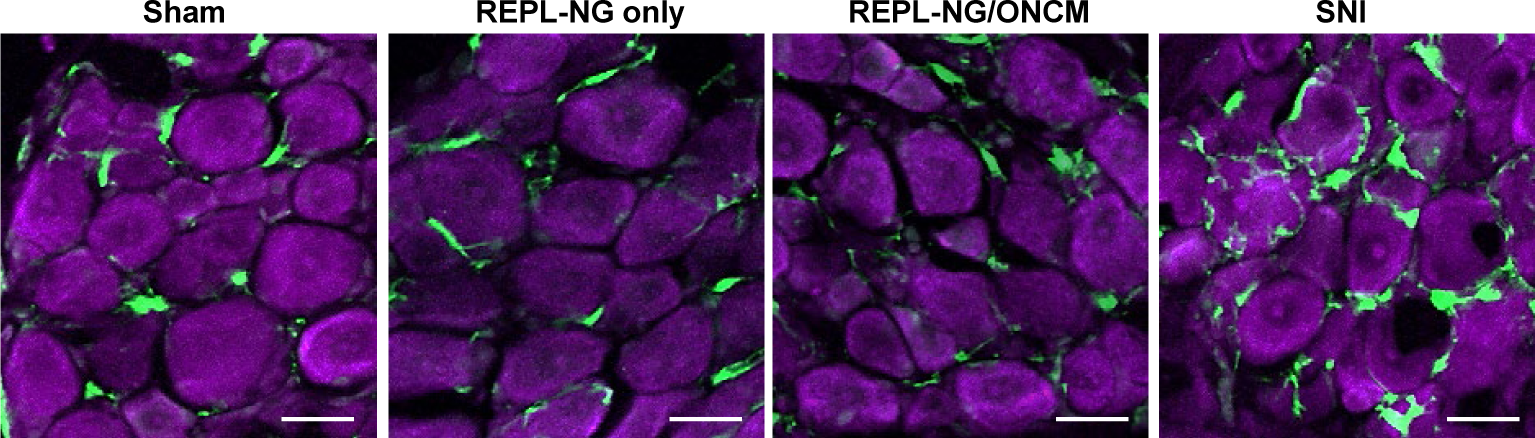
Intraganglionic injection of REPL-NG/ONCM does not induce activation of macrophages in DRGs. A, Representative images of DRG sections stained with Iba-1 (green) and NeuroTrace (violet) 14 d after intraganglionic injection of REPL only or REPL-NG/ONCM. DRGs were freshly dissected from animals at 0 (sham), 14 d after SNI (SNI). Scale bar, 50µm.

**Supplementary figure 8.**
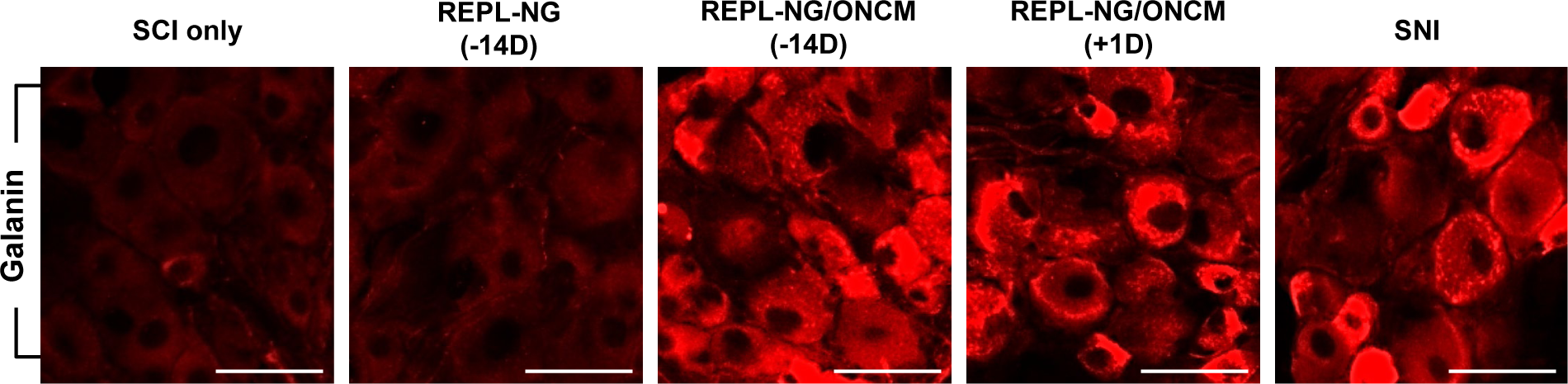
Intraganglionic injection of REPL-NG/ONCM upregulates galanin immunoreactivity. A, Representative images of Galanin-immunostained DRG sections from animals subjected to spinal cord injury (SCI) only or injected with REPL-NG 14 d before injury (REPL-NG; -14 D), with REPL-NG/ONCM 14 d before injury (REPL-NG/ONCM; -14 D), or with REPL-N G/ONCM 1 d after injury (REPL-NG-ONCM; +1D), and those subjected to preconditioning SNI before creating the spinal lesion (SNI). Scale bars, 100µm.

